# Rotavirus spike protein VP4 mediates viroplasm assembly by association to actin filaments

**DOI:** 10.1101/2022.06.08.495416

**Authors:** Janine Vetter, Guido Papa, Michael Seyffert, Kapila Gunasekera, Giuditta De Lorenzo, Mahesa Wiesendanger, Jean-Louis Reymond, Cornel Fraefel, Oscar R. Burrone, Catherine Eichwald

## Abstract

The formation of viroplasms is a well-conserved step in the rotavirus (RV) life cycle. In these structures, both virus genome replication and progeny assembly take place. A stabilized microtubule cytoskeleton and lipid droplets are required for the viroplasm formation, which involves several virus proteins. The viral spike protein VP4 has not previously been shown to have a direct role in viroplasm formation. However, it is involved with virus-cell attachment, endocytic internalization, and virion morphogenesis. Moreover, VP4 interacts with actin cytoskeleton components, mainly in processes involving virus entrance and egress, and thereby may have an indirect role in viroplasm formation. In this study, we used reverse genetics to construct a recombinant RV, rRV/VP4-BAP, which contains a biotin acceptor peptide (BAP) in the K145-G150 loop of the VP4 lectin domain, permitting live monitoring. The recombinant virus was replication competent but showed a reduced fitness. We demonstrate that rRV/VP4-BAP infection, as opposed to rRV/wt infection, did not lead to a reorganized actin cytoskeleton as viroplasms formed were insensitive to drugs that depolymerize actin and inhibit myosin. Moreover, wt VP4, but not VP4-BAP, appeared to associate with actin filaments. Similarly, VP4 in co-expression with NSP5 and NSP2 induced a significant increase in the number of viroplasm-like structures. Interestingly, a small peptide mimicking loop K145-G150 rescued the phenotype of rRV/VP4-BAP by increasing its ability to form viroplasms and hence, improve virus progeny formation. Collectively, these results provide a direct link between VP4 and the actin cytoskeleton to catalyze viroplasm assembly.

**IMPORTANCE:** The spike protein VP4 participates in diverse steps of the rotavirus (RV) life cycle, including virus-cell attachment, internalization, modulation of endocytosis, virion morphogenesis, and virus egress. Using reverse genetics, we constructed for the first time a recombinant RV, rRV/VP4-BAP, harboring a heterologous peptide in the lectin domain (loop K145-G150) of VP4. The rRV/VP4-BAP was replication-competent but with reduced fitness due to a defect in the ability to reorganize the actin cytoskeleton, which affected the efficiency of viroplasm assembly. This defect was rescued by adding a permeable small-peptide mimicking the wild-type VP4 loop K145-G150. In addition to revealing a new role of VP4, our findings suggest that rRV harboring an engineered VP4 could be used as a new dual vaccination platform providing immunity against RV and additional heterologous antigens.

## Introduction

Rotavirus (RV) is the primary etiological agent responsible for severe gastroenteritis and dehydration in infants and young children worldwide (1), resulting in the death of 128’000 children under five years of age in developing countries. Moreover, it also infects young animals such as piglets, calves, and poultry, negatively impacting the livestock industry (2-4).

RV virions are non-enveloped icosahedral particles composed of three concentric layers, called triple-layered particles (TLPs). The virus core-shell encloses eleven double-stranded (ds) RNA genome segments and twelve copies of both the RNA-dependent RNA polymerase, VP1, and the guanyl-methyltransferase, VP3 (5, 6). The icosahedral core-shell (T=1, symmetry) is composed of twelve decamers of VP2 and surrounded by 260 trimers of the structural protein VP6, constituting transcriptionally active double-layered particles (DLPs) (7, 8). Trimers of VP7 glycoprotein organized in an icosahedral symmetry (T=13) stand on the top of each VP6 trimer constituting the main building component of the virion outer layer. The spike protein VP4 anchors at each of the virion 5-fold axes adopting a trimeric conformation although having a dimeric appearance when visualized from the top of the capsid surface (9-12).

Immediately after RV internalization, the DLPs are released into the cytoplasm and become transcriptionally active (13), leading to the first round of transcription, which is necessary for halting the host innate immune response (14-18), shutting off the host translation machinery (19), and starting the building-up of viroplasms (20-22). The RV cytosolic replication compartments, termed viroplasms, are membrane-less electron-dense globular inclusions responsible for virus genome replication and virus progeny assembly (20, 23). The RV proteins NSP5, NSP2, and VP2 are the assembling blocks for viroplasms (24). Other virus proteins are also found in the viroplasms, including NSP4, VP1, VP3, and VP6, together with double-stranded and single-stranded viral RNAs. Host components, such as microtubules, lipid droplets, or miRNA-7 (25-27) are also recruited to viroplasms. The formation of viroplasms requires the reorganization and stabilization of the microtubule network and the recruitment of lipid droplets in a process directly associated with NSP2 and VP2. Thus, to be formed, the viroplasms have to coalesce from small-punctuated structures diffused in the cytosol at early times of infection to perinuclear large-mature viroplasms at late times of infection (25, 26, 28). However, pieces of evidence suggest that not only microtubules but also actin and intermediate filaments reorganize under RV infection by using a mechanism not yet described (25, 29). Many virus proteins have multifunctional roles during the viral life cycle, and the RV proteins are no exception. An example is VP4 which is cleaved by a trypsin-like enzyme found in the intestinal tract (30) into two main products, VP8* (28 kDa, amino acids 1-247) and VP5* (60 kDa, amino acids 248-776) that remain non-covalently associated with the infectious RV virion allowing the initialization of the virus entry process (31, 32). VP8* and a significant portion of VP5*, VP5CT (amino acids 246-477), constitute the distal globular and the central body of the spike, respectively (33, 34). The VP8* subunit has several functions, such as haemagglutinin activity (35), involvement in binding the host sialic acid (34), and a determinant role in virus tropism. VP5* has been implicated in the interaction with integrins (36-38). During virus internalization, VP4 engages the endocytic pathway (39, 40) by binding to the small GTPase Rab5 and PRA1 within early endosomes (41), directly activating hsp70 (42, 43) and associating with the actin-binding protein Drebrin 1, an RV restriction factor (44). VP4 also plays an essential role in virion morphogenesis and can be found as a soluble protein in the cytosol (45). When expressed in the absence of other RV proteins, VP4 is associated with the microtubules and the actin cytoskeleton (45-50). In polarized cells, VP4 seems to interact with the apical actin cortex, leading to the remodeling of the brush border and subsequently releasing the RV virions into the medium (48). This actin association is dependent on a VP4 actin-binding domain present at its C-terminus (residues 713-776) in cooperation with the coiled-coil domain (residues 481-574) (45). However, there is no direct evidence that VP4 participates in actin reorganization during RV infection.

Here, we describe the generation of a recombinant RV (rRV) harboring a genetically modified genome segment 4 (gs4) encoding the spike protein VP4 with a <u>b</u>iotin-<u>a</u>cceptor <u>p</u>eptide (BAP) tag of 15 amino acids inserted in an exposed loop of VP8* (residues K145-G150). Even though rRV/VP4-BAP is internalized and able to produce virus progeny, it has significantly reduced virus fitness because of an impaired ability of VP4-BAP to associate with the actin cytoskeleton. In addition, we provide clear evidence that VP4 acts as a catalyst for the assembly of the viroplasms in an actin-dependent process.

## Results

### Production and analysis of recombinant rotavirus harboring a BAP tagged spike protein

As the VP4 spike protein plays an essential role in the host cell tropism, attachment, and internalization, we addressed if VP4 could be engineered by incorporating a heterologous peptidic tag within its coding sequence without compromising its structural and functional properties. To test this hypothesis, we identified in the previously published crystal structure of simian RRV VP4 (10) four different exposed loops localized in the lectin domain (amino acids 65-224) of the VP8* subunit and then inserted a <u>b</u>iotin <u>a</u>cceptor <u>p</u>eptide (BAP) tag (51, 52) in each of the corresponding loops of the VP4 simian strain SA11.

As depicted in **Fig. 1a**, the selected loops for the BAP tag insertions were T96-R101, E109-S114, N132-Q137, and K145-G150, labeled with the colors blue, orange, pink, and green, respectively. First, we assessed the biotinylation of these four BAP-tagged VP4 proteins, herein VP4-BAP, in cell lysates by western blot-retardation assay (WB-ra) (51). For this purpose, each construct was co-expressed with cytosolically localized biotin ligase Bir A (cyt-BirA) in MA104 cells infected with recombinant vaccinia virus encoding T_7_ RNA polymerase to allow the synthesis of cytosolic VP4 transcripts. As the synthesis of nuclear VP4 transcripts promotes undesired mRNA splicing (53), we used a T_7_ promoter to favor cytosolic transcription provided by the T_7_ RNA polymerase. As shown in **Fig. 1b**, the four VP4-BAP variants (**Fig. 1b**, lanes 4, 6, 8 and 10), but not the wild type (wt) VP4 (**Fig. 1b**, lane 2), were fully biotinylated when the cell extracts were incubated with streptavidin (StAv), generating a band shift of approximately 140 kDa that corresponds to the VP4-BAP/StAv complex. Of note, biotinylated protein-StAV complexes hinder epitopes of the biotinylated protein denoted as a shifted and weaker band in western blot. The band observed immediately above the VP4-BAP band (80 kDa) represents a phosphorylated form of the protein that is only present in transfected cells but not in RV-infected cells, as demonstrated by the λ-phosphatase treatment of the cellular extracts (Fig. S1a and b). This result suggests that the expression and stability of the different VP4-BAP proteins were not affected by the location of the inserted BAP tag.

**Figure 1.**
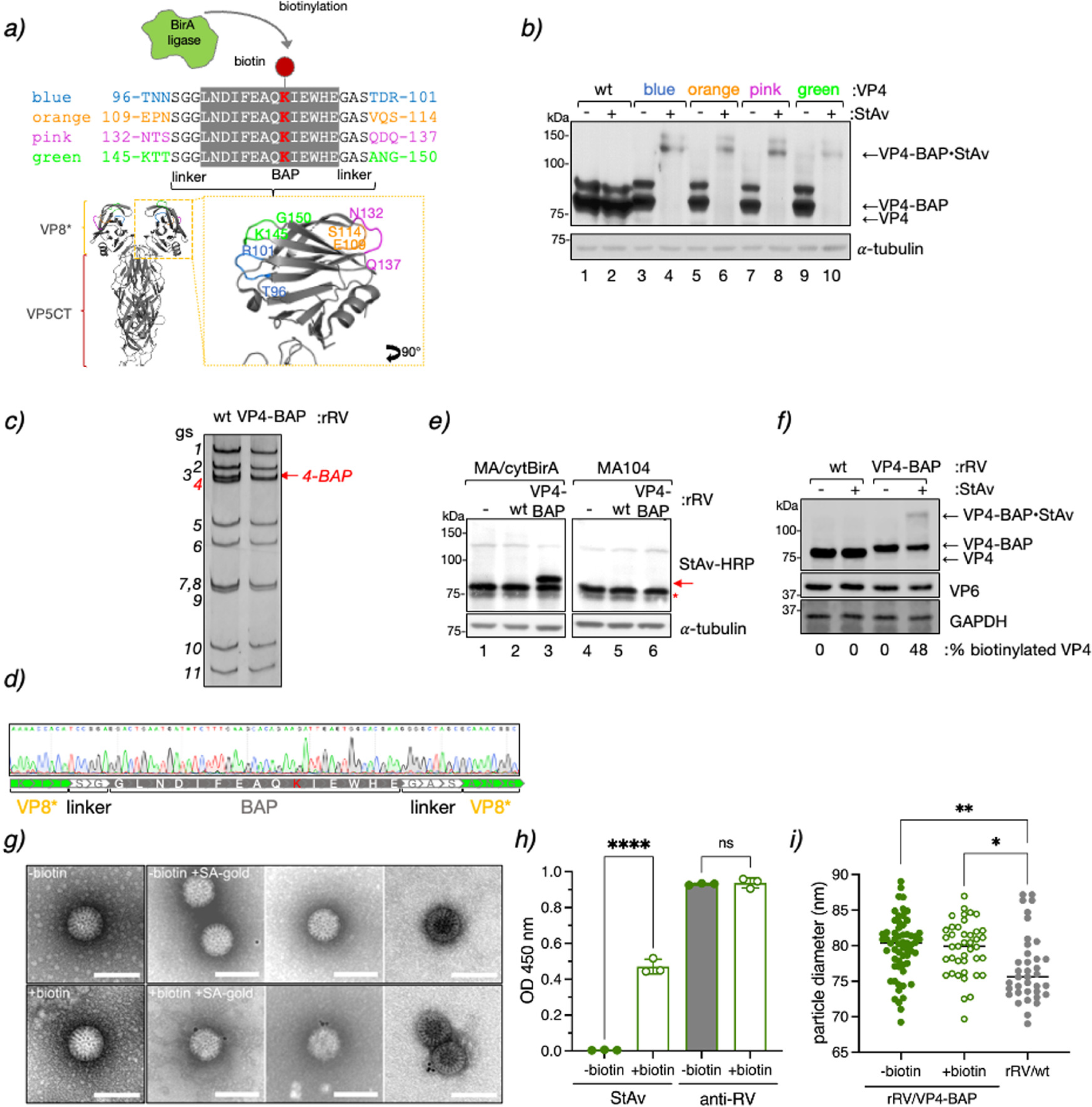
Generation of VP4-BAP tagged recombinant rotavirus. **(a)** Schematic representation of BAP tag inserted into the lectin domain loops of the VP8* subunit of VP4 from RV simian strain SA11 (GenBank: X14204.1). The lysine (K, red) indicates the biotinylation site of BirA ligase. Four different VP4 proteins tagged with BAP (VP4-BAP) were built between amino acid regions T96-R101 (blue), E109-S114 (orange), N132-Q137 (pink), and K145-G150 (green). VP4 trimer ribbon structure is presented for the visualization of VP8* (yellow) and VP5CT (body and stalk, red) fragments. An inset in VP8* indicates the different positions in the hydrophobic loops of VP8* where the BAP tags were inserted and colored in blue, orange, pink, and green. **(b)** Immunoblot retardation assay of cell lysates transiently expressing wtVP4 and VP4-BAP, tagged at blue, orange, pink, and green positions, respectively. Untreated (-) and streptavidin-treated (+) samples are indicated. The membrane was incubated with anti-VP4 to detect unbound VP4-BAP and VP4-BAP bound to streptavidin (VP4-BAP•StAv). α-tubulin was used as a loading control**. (c)** dsRNA electropherotype of the genome segments of rRV/wt and rRV/VP4-BAP. The red arrow points to gs4-BAP. **(d)** Sequence chromatogram of gs4 of rRV/VP4-BAP. The sequence indicates the position of the linkers and the BAP tag in between the VP8*. **(e)** Immunoblotting of uninfected (lanes 1 and 4) or infected cell lysates in MA-cytBirA cells (left panel) or MA104 cells (right panel) infected with either rRV/wt (lanes 2 and 5) or rRV/VP4-BAP (lanes 3 and 6) [MOI, 25 VFU/cell]. Biotinylated proteins were detected with StAv-HRP. α-tubulin was used as a loading control. The red arrow and red star indicate biotinylated VP4-BAP and host undetermined biotinylated protein, respectively. **(f)** Immunoblot retardation assay of MA-cytBirA cell lysates infected with rRV/wt or rRV/VP4-BAP untreated or treated with StAv. The membrane was incubated with anti-VP4 and anti-VP6 for detection of the virus. GAPDH was used as a loading control. The percentage of biotinylated VP4 and VP4-BAP was normalized to VP6 expression. **(g)** Visualization at a high resolution of purified virions isolated of rRV/VP4-BAP infected MA-cytBirA cells untreated (-biotin, upper panel) or treated (+biotin, lower panel) with 100µM biotin. After purification, the virions were labeled with streptavidin conjugated to colloidal gold (12 nm), followed by negative staining electron microscopy (right panel). The scale bar is 100 nm. **(h)** Detection of purified unbiotinylated (grey bars) and biotinylated (open bars) rRV/VP4-BAP particles. The particles were coated in an ELISA plate followed by binding to streptavidin-HRP or guinea pig anti-rotavirus followed by anti-guinea pig conjugated to HRP. The median from three independent experiments is shown (ordinary one-way ANOVA, (****) p-value<0.0001). **(i)** Scatter dot plot comparing the diameter of purified particles from unbiotinylated (-biotin, filled green dots) or biotinylated (+biotin, open green dots) RV/VP4-BAP and rRV/wt (filled grey dots). The median value is indicated, n>40 particles, ordinary one-way ANOVA, (*) p-value<0.05 and (**) p-value<0.01.

Next, we assessed whether the four VP4-BAP proteins could assemble into infectious RV particles and support virus replication. To rescue recombinant rotavirus (rRV) harboring a genetically modified genome segment 4 (gs4) encoding for the different VP4-BAP proteins (gs4-BAP), we took advantage of the previously established reverse genetics system (54). Of the four different constructs, we successfully rescued only the rRV harboring gs4-BAP encoding the BAP tag within the K145-G150 loop in VP8* (**Fig. 1a**), herein named rRV/VP4-BAP. The rescued virus was confirmed by the differential migration pattern of the modified gs4-BAP compared to the wt gs4 (**Fig. 1c**) and by Sanger sequencing (**Fig. 1d**). Notably, the genome segment 4 (gs4-BAP) and all the other ten genome segments of rRV/VP4 were stable in tissue culture at least until virus passage ten as determined by Sanger sequencing and deep sequencing (Fig S1b and supplementary information). We then investigated the ability of rRV/VP4-BAP to express biotinylated VP4-BAP. Specifically, cell extracts of rRV/VP4-BAP infected MA/cytBirA cells (MA104 cells stably expressing cytosolic localized BirA) were analyzed at 6 hpi by western blot. Thus, the produced VP4-BAP protein showed to be biotinylated as demonstrated after incubation with StAv-peroxidase, which detected a band of approx. 85 kDa only in rRV/VP4-BAP infected MA/cytBirA cell extracts but not in rRV/VP4-BAP-infected MA104 cells (**Fig. 1e**, lane 3 and 6). As expected, VP4 biotinylation was detected neither in rRV/wt infected MA/cytBirA cells nor in rRV/VP4-BAP infected MA104 cells (**Fig. 1e**, lanes 2, 5, and 6). We found by WB-ra that the fraction of biotinylated VP4-BAP corresponds to 48 % of the total protein (**Fig. 1f**).

We next examined if the biotinylated virus-encoded VP4-BAP is incorporated into newly assembled virus particles. For this, we purified rRV/VP4-BAP virions produced in MA/cytBirA cells in the absence or presence of biotin and visualized the virus particles by negative staining electron microscopy after incubation with StAv conjugated to gold particles. Thus, 53% of the virions produced in the presence of biotin were decorated with StAV-gold particles (**Fig. 1g**) but none in the unbiotinylated control particles. Similarly, indirect ELISA with identical amounts of unbiotinylated and biotinylated purified rRV/VP4-BAP revealed a signal upon StAV-peroxidase staining only for biotinylated samples. Furthermore, similar signals were observed in ELISA using an anti-rotavirus antibody (**Fig. 1h**). These outcomes collectively suggest that virus-encoded VP4-BAP is biotinylated and incorporated in newly formed RV particles. Interestingly, as shown in **Fig. 1i**, the rRV/VP4 particles appear to have a slightly larger diameter (∼80 nm) when compared to rRV/wt particles (∼75 nm) but were still in the range of TLPs (55).

### Subcellular localization of rRV/VP4-BAP

We investigated the subcellular localization of the newly produced biotinylated VP4-BAP in rRV/VP4-BAP infected cells at 6 hpi, a time point showing well-assembled viroplasms (25). For this, both rRV/wt and rRV/VP4-BAP-infected MA/cytBirA cells were incubated in the absence or presence of biotin for 4 hours before fixation. Biotinylated VP4-BAP detected with StAv-Alexa 555 was found surrounding the viroplasms (stained anti-NSP5) (**Fig. 2a**). However, no StAv-Alexa 555 signal was detected in cells infected with either rRV/wt or rRV/VP4-BAP without biotin. Notably, the biotinylated VP4-BAP partially co-localized with trimeric VP7 in the endoplasmic reticulum (ER) (**Fig. 2b**). Additionally, the cytosolic distribution of VP4-BAP was similar to that of VP4 in rRV/wt-infect cells **(Fig. 2c)**. These results suggest that the modification exerted in VP4-BAP does not impact VP4 subcellular localization in infected cells.

**Figure 2.**
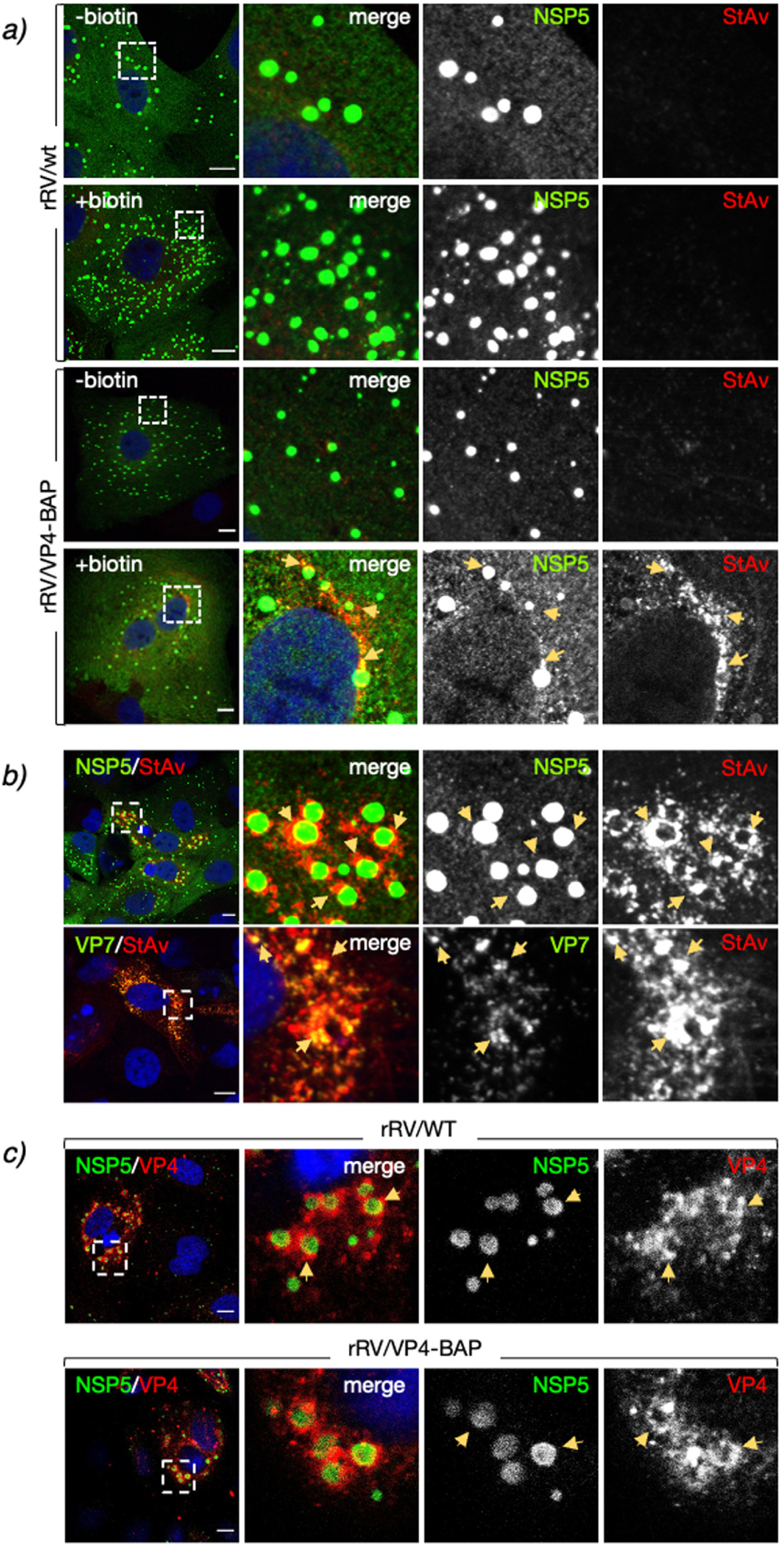
RV protein localization upon rRV/VP4-BAP infection. **(a)** rRV/wt (top panel) and rRV/VP4-BAP (bottom panel) infected MA-cytBirA cells were untreated (-biotin) or treated with biotin (+biotin). Cells were fixed with paraformaldehyde at 6 hpi and stained for the detection of viroplasms (anti-NSP5, green) and biotinylated proteins (streptavidin-Alexa 555, red). Nuclei were stained with DAPI (blue). **(b)** Immunostaining images of rRV/VP4-BAP infected MA-cytBirA cells in the presence of biotin. At 6 hpi, paraformaldehyde-fixed cells were stained for the detection of VP4-BAP (StAv, red) in viroplasms (anti-NSP5, green) (upper row) or mature RV particles (anti-VP7 clone 159, green) (bottom row). **(c)** Immunofluorescence images of cells infected with rRV/wt (upper row) or rRV/VP4-BAP (lower row) comparing the localization of VP4 and VP4-BAP (anti-VP4, red), respectively. The viroplasms were detected with anti-NSP5 (green). Nuclei were stained with DAPI. In all the figures, the dashed white boxes correspond to the image insets of the right columns. The yellow arrows point to the VP4-BAP or VP4 signal. The scale bar is 10 µm.

### Impaired virus fitness of rRV/VP4-BAP

To compare the replication fitness of rRV/VP4-BAP with that of rRV/wt, we infected MA104 cells at equal MOI and harvested the virus at various time points until 48 hpi. As depicted in **Fig 3a**, rRV/VP4-BAP showed a significantly delayed fitness curve compared to rRV/wt. To investigate this divergence in the virus replication, we infected cells with identical MOI (**Fig. 3b**) or an equal number of virus particles (**Fig. 3c**) and quantified cells showing viroplasms at 6 hpi. In both experimental conditions, we observed a significantly reduced ratio of cells containing viroplasms upon rRV/VP4-BAP infection when compared with rRV/wt infection.

**Figure 3.**
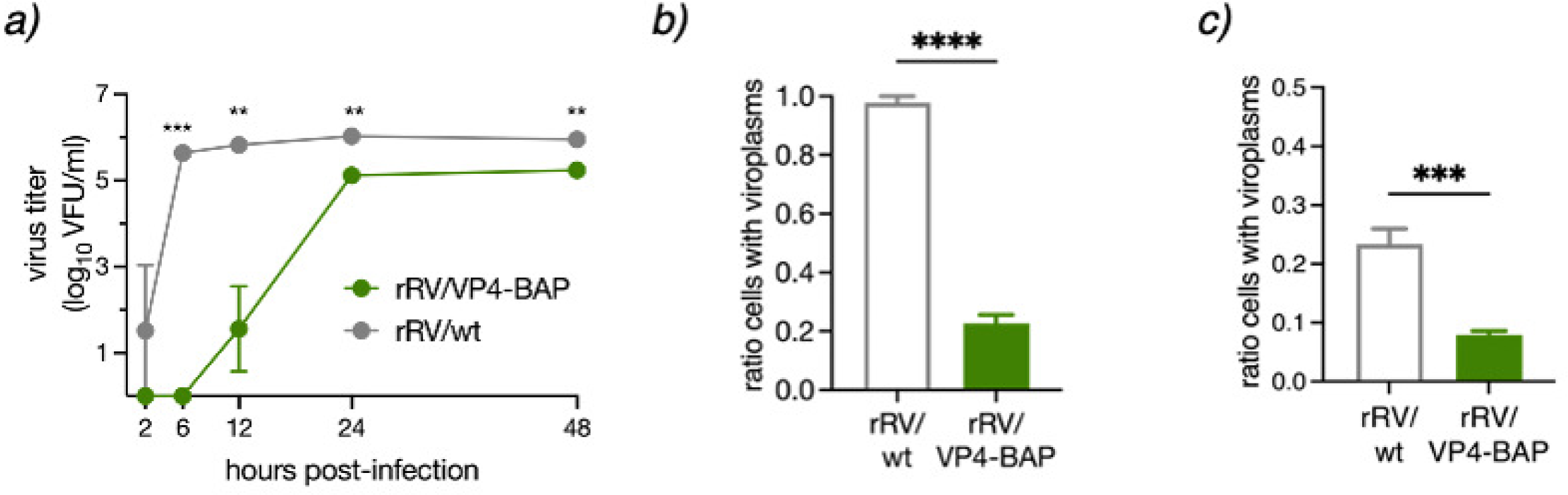
Replication fitness of rRV/VP4-BAP is delayed. **(a)** Virus replication fitness curve from 0 to 48 hpi of rRV/wt and rRV/VP4-BAP. The curve represents the mean of three independent experiments. Welch’s t-test, (*), p<0.05; (***), p<0.001 and ns, not significant. Quantification of cells with viroplasms of MA104 cells infected rRV/wt and rRV/VP4-BAP at **(b)** same MOI (25 VFU/cell) or **(c)** the same numbers of virus particles.

### Comparable entry processes between rRV/wt and rRV/VP4-BAP

Since VP4 has an essential role in virus-cell attachment, we interrogated whether the ability of rRV/VP4-BAP particles to bind to cells was impaired. To test this hypothesis, we performed a nonradioactive binding assay described previously by Zarate *et al.* (56) to compare the attachment to MA104 cells with different amounts of either rRV/wt or rRV/VP4-BAP. As depicted in **Fig 4a**, no differences in cell attachment were observed among the two viruses. Of note, the antibody conditions used for the virus detection are in the linear range (**Fig S2a**). Moreover, biotin-labeled rRV/VP4 did not hinder the virus-cell attachment as denoted by the same results for the attachment of both unbiotinylated and biotinylated rRV/VP4-BAP virus particles (**Fig 4b**). As expected, binding of rRV/VP4-BAP was detected with StAVonly when grown in the presence of biotin (**Fig 4c**).

**Figure 4.**
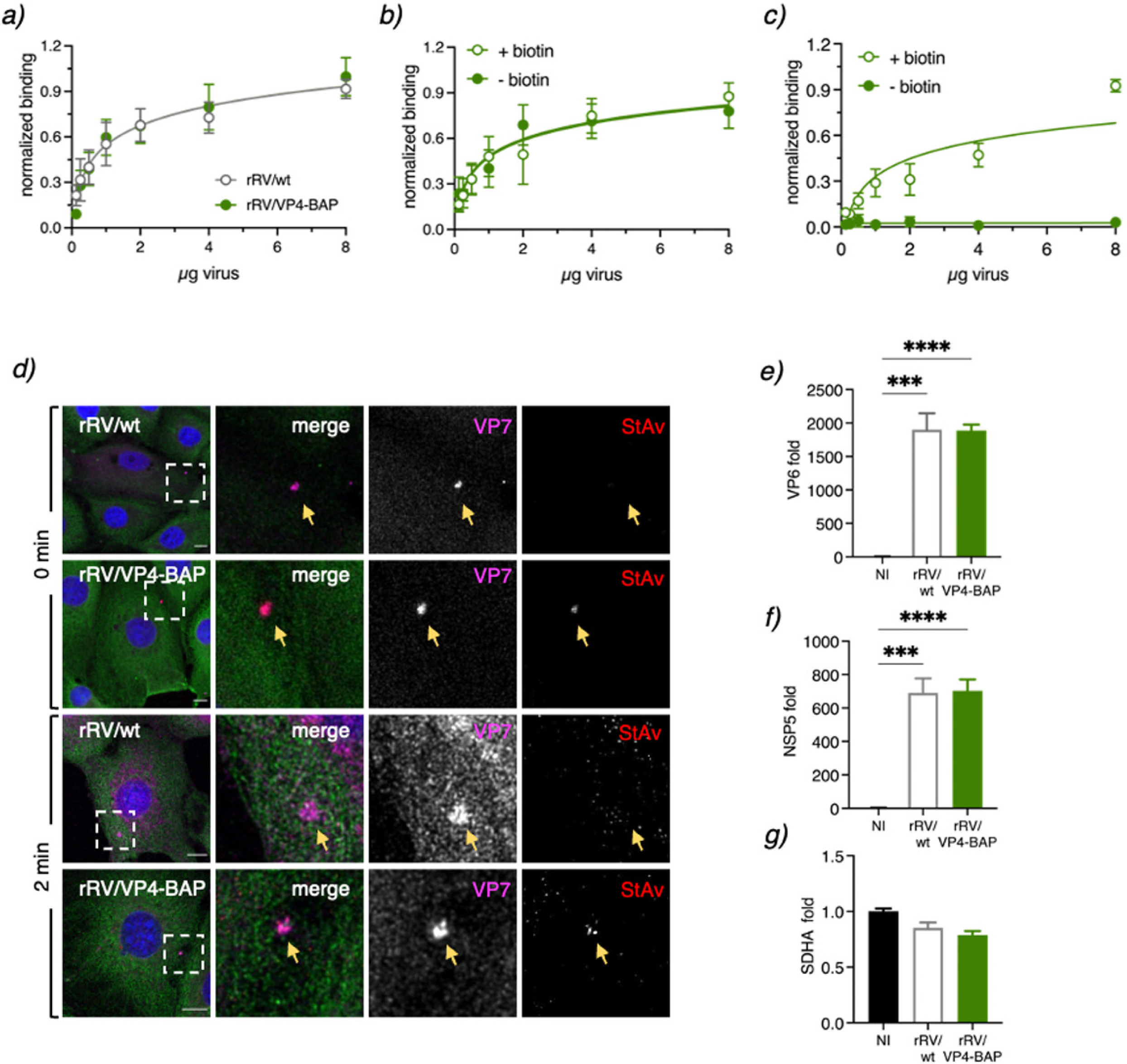
Virus attachment and internalization of RV/wt and rRV/VP4-BAP are comparable. **(a)** Virus attachment to MA104 cell surface with different amounts of purified rRV/wt (open grey dots) and rRV/VP4-BAP (full green dots). Samples were detected using a guinea pig anti-RV. The result corresponds to the mean of nine normalized independent experiments. Data analysis corresponds to nonlinear regression, p=0.2987, n=9. Virus attachment assay to MA104 cell surface with different amounts of unbiotinylated (-biotin, full green dots) and biotinylated (+biotin, green open dots) purified rRV/VP4-BAP detected with **(b)** guinea pig anti-RV followed by secondary antibody conjugated to HRP. Data analysis corresponds to nonlinear regression, p=0.6151, n=6. **(c)** Same as above, but samples were detected with streptavidin-HRP. Data analysis corresponds to nonlinear regression, p<0.0001, n=6. **(d)** Internalization of purified virions into MA104 cells at 0 min (upper panel) and 2 min (lower panel). Purified virions of rRV/wt and biotinylated rRV/VP4-BAP were previously labeled with StAv-Alexa 555 (red). At the indicated time post-infection, cells were fixed and immunostained for VP7 trimers (mAb anti-VP7 clone 159, pink) and MTs (anti-α-tubulin, green). Nuclei were stained with DAPI (blue). White open boxes indicate the magnified images at the right. Arrows point to virus particle clumps detected by anti-VP7. The scale bar is 20 µm. **(e-g)** Plots from quantitative RT-PCR comparing rotavirus transcripts of NSP5, VP6, and housekeeping gene (SDHA) in non-infected (NI), rRV/wt- and rRV/VP4-BAP-infected cellular extracts at 4 hpi. The results correspond to the mean ± SEM of three independent experiments, ordinary one-way ANOVA, (**), p<0.01; (***), p<0.001; (****), p<0.0001.

Next, we investigated whether the delay in viral replication fitness was caused by a difference in virus internalization. Purified rRV/VP4-BAP and rRV/wt virions were compared and analyzed for virus internalization by confocal scanning laser microscopy (CSLM) using immunostaining with the conformational monoclonal antibody anti-VP7 (clone 159), which only recognizes the trimeric form of the VP7 protein (57, 58). As a control, purified rRV/VP4-BAP, but not rRV/wt, was directly labeled with StrAv-Alexa 555 prior to infection. Initially (0 min), VP4-BAP and VP7 signals co-localized on the cell surface, indicating the association of virions to the cell membrane. However, after two minutes at 37°C, both signals were already internalized (**Fig 4d**). The localization patterns were comparable to those observed at the same time points with rRV/wt virions, suggesting no differences in the internalization mechanism between the two viruses.

Since virus-cell attachment and virus internalization were comparable between rRV/wt and rRV/VP4, we investigated if virus transcription was defective or delayed for rRV/VP4-BAP. Thus, we compared the abundance of NSP5 and VP6 virus transcripts at 4 hpi of MA104 cells infected with either rRV/wt or rRV/VP4-BAP. As denoted in **Figs 4e-g**, the transcription levels of NSP5, VP6, and the housekeeping gene SDHA were comparable in cells infected with rRV/wt or rRV/VP4-BAP.

### rRV/VP4-BAP has a defect in a step between virus transcription and viroplasm formation

We analyzed by high-definition electron microscopy the structural morphology of the viroplasms from rRV/wt or rRV/VP4-BAP at two time points, 6 and 12 hpi, which for simian strain SA11 corresponds to a time showing well-formed viroplasms and a time with highly mature viroplasms, respectively (**Fig 5a**). No apparent differences in the viroplasm morphology were observed between the two viruses at 6 and 12 hpi. Similarly, we examined whether the liquid-liquid phase separation properties of rRV/VP4-BAP viroplasms were modified. For this, we took advantage of our previously established MA104 cell line stably expressing NSP2-mCherry (MA/NSP2-mCherry) to visualize viroplasm formation in living cells because of the ability of this protein to get recruited into viroplasms during RV infection (21, 25, 59). Next, we measured the NSP2-mCherry diffusion dynamics in single viroplasms using fluorescence recovery after photobleaching (FRAP) experiments (**Fig 5b and c**). We found that FRAP properties concerning NSP2-mCherry half-time recovery and mobility were similar for both viruses (**Fig. 5d and e**).

**Figure 5.**
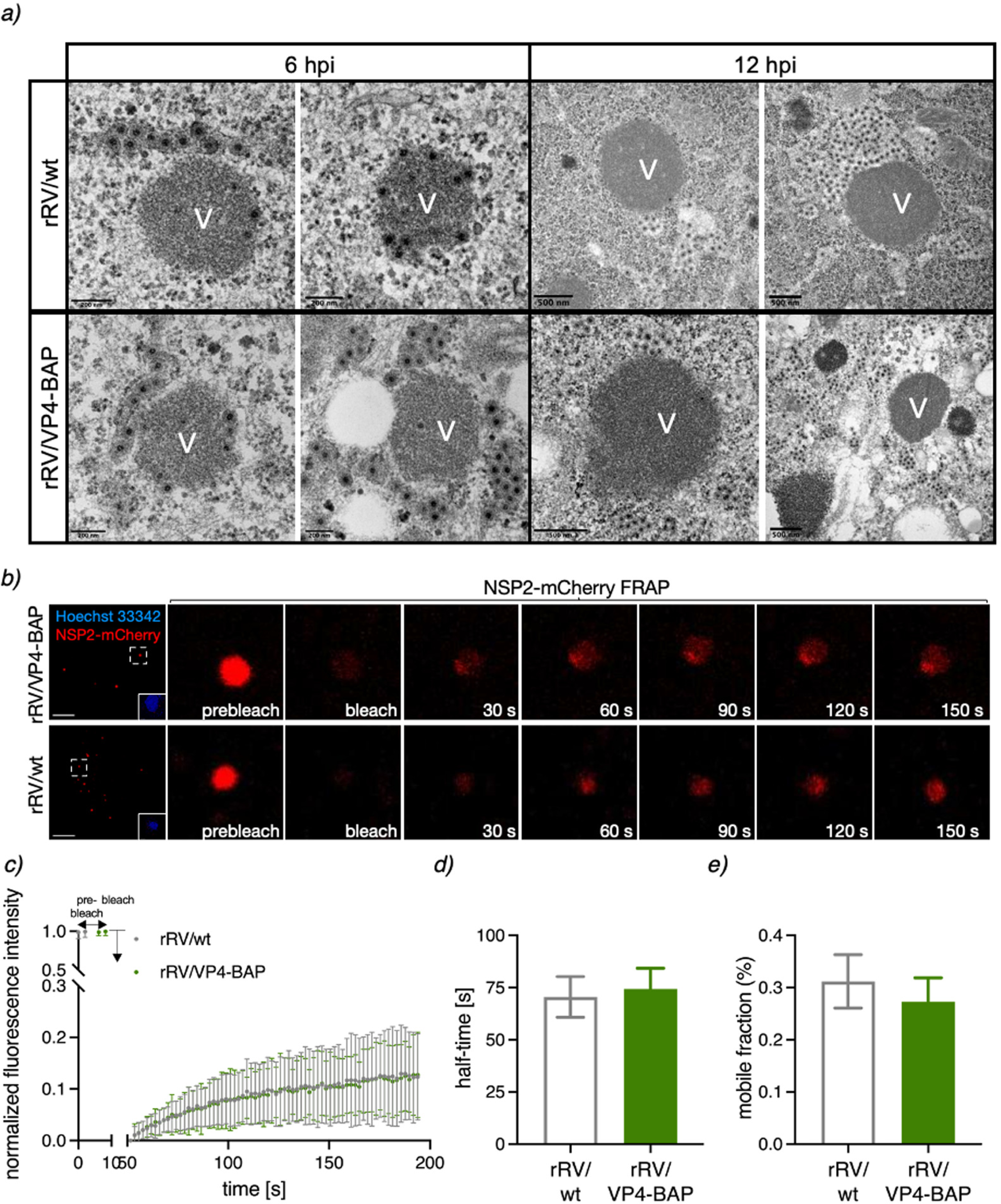
Viroplasm morphology and behavior for rRV/wt and rRV/VP4-BAP. **(a)** Representative high-resolution electron micrograph of viroplasms from rRV/wt (upper row) and rRV/VP4-BAP (lower row) infected MA104 cells at 6 hpi (left panel) and 12 hpi (right panel). Scale bars are 200nm and 500 nm, as indicated. **(b)** Fluorescence images of FRAP measurement of single viroplasm of cells infected with rRV/VP4-BAP (top) and rRV/wt (bottom) at pre-bleach, post-bleach, and recovery time conditions. Each inset indicates the bleached viroplasm of the images at the right. Nuclei were stained with Hoechst 33342. The scale bar is 10 µm. **(c)** FRAP recovery curve of NSP2-mCherry in single viroplasm from rRV/VP4-BAP (green) and rRV/wt (grey) infected MA/NSP2-mCherry cells at 5 hpi (n=27 and 25, respectively). Plots indicating recovery half-time **(d)** and diffusion **(e)** means of NSP2-mCherry in single viroplasms of rRV/VP4-BAP and rRV/wt after photobleaching.

To confirm our results and exclude the involvement of the endocytic pathway, we purified and transfected MA104 cells with an equal number of DLPs of the two viruses (**Fig 6a**). Of note, purified rRV/wt DLPs or rRV/VP4-BAP DLPs had the same size, while rRV/VP4-BAP TLPs were larger than rRV/wt TLPs (**Fig 1i and 6a**). At 6 h after transfection, we observed identical expression of NSP5 for both viruses, as denoted by in-cell western assay **(Figs 6b and c)**. In the same experimental setting, the number of cells showing viroplasms (detected with anti-NSP5) was significantly reduced in rRV/VP4-BAP infected cells compared to rRV/wt infected cells (**Figs 6d and e**). These outcomes strongly suggest that the reduced replication fitness of rRV/VP4-BAP involves a step in viroplasm assembly. **VP4 promotes viroplasm assembly.** Since we hypothesized that VP4 may have a yet unidentified role in the viroplasm assembly, we investigated if spontaneous disruption of the viroplasm led to a delayed re-assembly of these structures. To challenge this hypothesis, we used 1,6-hexanediol (1,6-HD), a well-described aliphatic alcohol able to disrupt key drivers of liquid-liquid phase separation and recently shown to be effective in dissolving RV viroplasms (28) and determined the recovery of NSP2-mCherry in viroplasms. For this, MA/NSP2-mCherry cells at 5 h after infection with either rRV/VP4-BAP or rRV/wt were treated for 6 min with 1,6-HD and observed for 30 min after washout of the compound (**Fig. 7a and b**). Interestingly, the recovery kinetic of rRV/VP4-BAP viroplasms was delayed compared to that of rRV/wt at short times after removing the drug (2 min), as confirmed by both a reduced ratio of cells presenting viroplasms (**Fig. 7c**) and a reduced number of viroplasm per cell (**Fig. 7d**). In order to further characterize the relationship between viroplasms and VP4, we performed transcriptional silencing experiments with VP4 specific siRNAs (siVP4) in MA/NSP2-mCherry cells (**Fig. 7e**). In this context, it has been previously described that the silencing of VP4 during RV infection leads to viroplasm formation even if TLPs assembly is impaired (60). We then monitored viroplasm formation in siVP4 and control-siRNA (scr) transfected MA104 cells infected with simian strain SA11 and treated with 1,6-HD as described for the experiment shown in **Fig 7a**. The viroplasms produced on VP4 silenced cells had a delayed recovery kinetic similarly to that observed for rRV/VP4-BAP viroplasms (**Fig 7f and g**). Furthermore, similarly to **Fig 7d**, the number of viroplasms per cell was significantly decreased in siVP4 treated cells at 2 min of recovery compared to the experimental controls (**Fig 7h**). Thus, these results strongly suggest that VP4 promotes either indirectly or directly the assembly of viroplasms.

**Figure 6.**
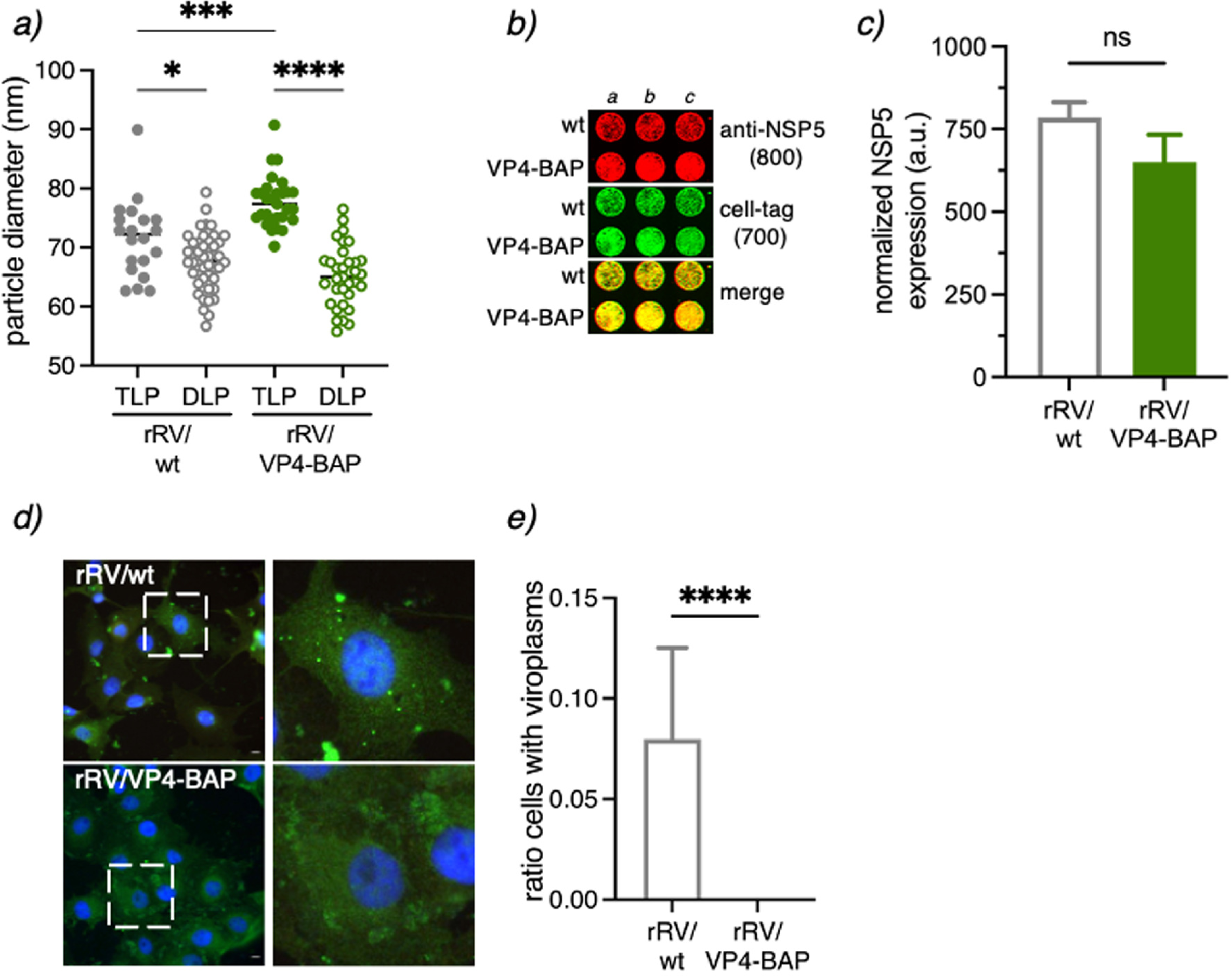
Transfection of rRV/VP4-BAP DLPs is inefficient in forming viroplasms. **(a)** Size plot of TLPs and DLPs purified from rRV/wt and rRV/VP4-BAP. The median value is indicated, n>30 particles, ordinary one-way ANOVA, (*) p-value<0.05 and (****) p-value<0.0001. **(b)** In-cell western blot of MA104 cells transfected for 4 h with identical numbers of rRV/wt and rRV/VP4-BAP DLPs. RV infection was detected by staining with anti-NSP5 (IDye800, red). The loading control corresponds to cell tag 700 (green). The merge of the two channels is shown at the bottom. **(c)** Plot showing normalized amounts of NSP5 in rRV/wt and rRV/VP4-BAP-DLPs transfected MA104 cells. Each experiment was done in triplicate. **(d)** Micrograph of rRV/wt and rRV/VP4-BAP DLPs transfected MA104 cells immunostained to detect viroplasms with anti-NSP5 (green). Nuclei were stained with DAPI (blue). Open dashed white box shows enlarged viroplasm inset at the right. The scale bar is 10 µm. **(e)** Quantification of the cells showing viroplasms upon transfection with DLPs of rRV/wt and rRV/VP4-BAP. The data correspond to the mean ± SD, Welch’s t-test, (****), p>0.0001.

**Figure 7.**
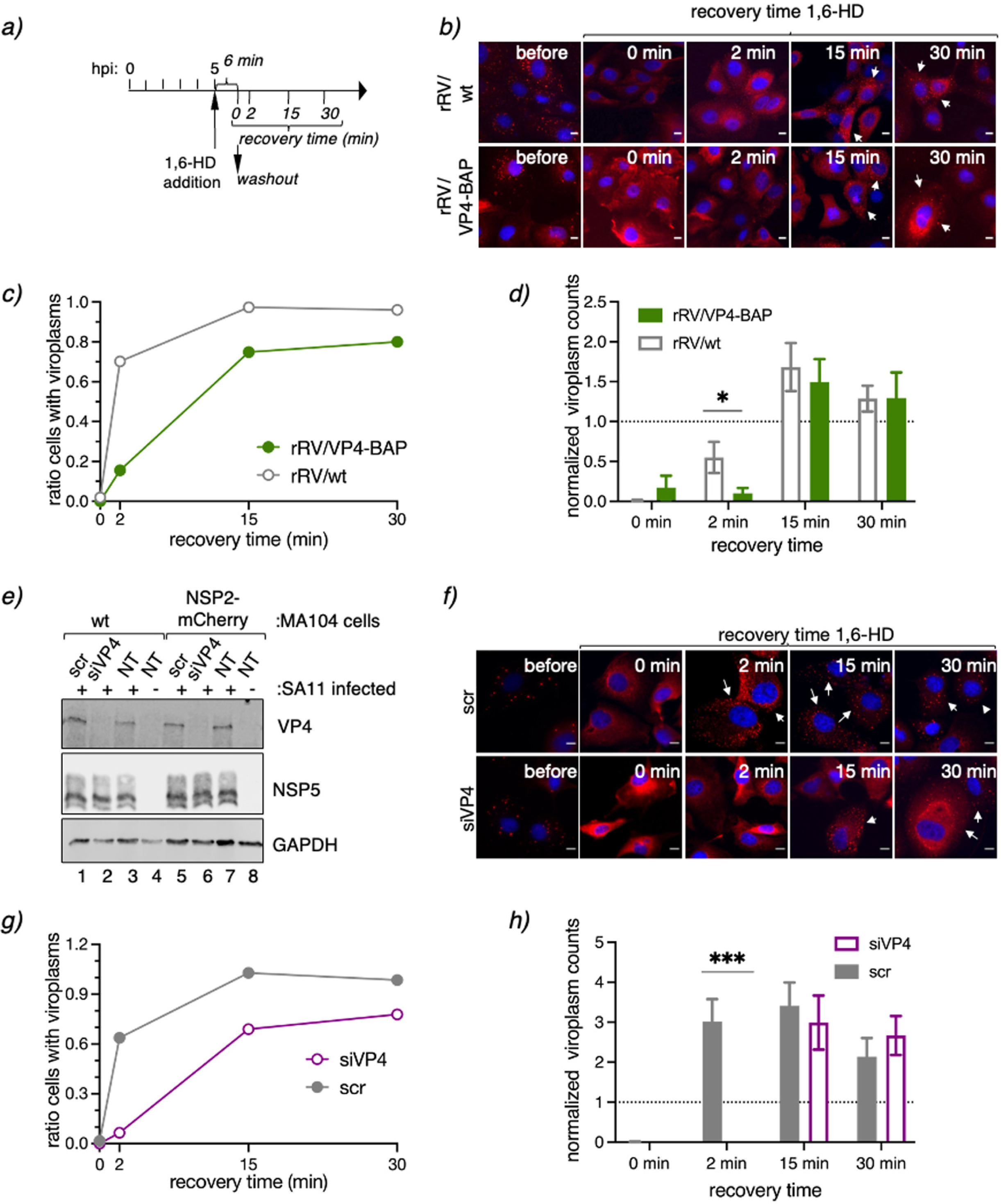
VP4 has a role in viroplasm formation. **(a)** Schematic representation for the characterization of LLPS condensates in viroplasms of RV-infected cells. At 5 hpi, RV-infected MA/NSP2-mCherry cells were treated with 1,6-HD for 6 min. The drug was washed out, and samples were fixed and imaged for viroplasm quantification at 0-, 2-, 15- and 30-min post-recovery. **(b)** Representative images of MA/NSP2-mCherry cells infected at 5 hpi with rRV/wt (upper row) or rRV/VP4-BAP (lower row) and treated with 3.5% of 1,6-HD. Cells were washed and monitored for viroplasm formation at 0-, 2-, 15- and 30-min post-recovery. White arrows point to cells with recovered viroplasms. The scale bar is 10 µm. Plots indicating the ratio of cells with viroplasms (**c**) and viroplasm counts per cell (**d**) normalized at initial conditions (5 hpi). **(e)** Immunoblot of cellular extracts prepared at 6 hpi from non-infected or SA11-infected MA104 or MA/NSP2-mCherry cells silenced with siVP4 or control siRNA (scr). The membrane was stained with anti-VP4, anti-NSP5, and anti-GAPDH (loading control). **(f)** Representative images of SA11-infected MA/NSP2-mCherry cells knocked down with scr (upper row) or siVP4 (lower row) and treated for 6 min with 3.5% of 1,6-HD. Cells were washed and monitored for viroplasm formation at 0-, 2-, 15- and 30-min post-recovery. White arrows point to cells showing recovered viroplasms. The scale bar is 10 µm. Plot indicating the ratio of cells with viroplasms (**g**) and viroplasm counts per cell (**h**) normalized to initial conditions after recovery from 1,6-HD treatment of SA11-infected MA104 cells silenced with siVP4 or scr The data represent the mean ± SEM Student’s t-test (*), p<0.05; and (***), p<0.001.

### VP4-BAP impairs VLS assembly and association to actin filaments

Viroplasm-like structures (VLS) are simplified models for the study of complex viroplasm structures, requiring the co-expression of NSP5 with either NSP2 (24) or VP2 (22) to form globular cytosolic inclusions morphologically similar to viroplasms but unable to replicate and produce virus progeny. Considering this rationale, we formed VLSs by expressing NSP5 and NSP2 in the presence of GFP, VP4-GFP, or VP4-BAP-GFP (**Fig 8a**). We noticed the absence of co-localization of either VP4-GFP or VP4-BAP-GFP in VLSs. Furthermore, when we quantified the number of VLSs per cell (**Fig 8b**), we noticed that the number of VLSs formed in the presence of VP4-GFP was much larger than that produced in the presence of VP4-BAP-GFP or GFP.

**Figure 8.**
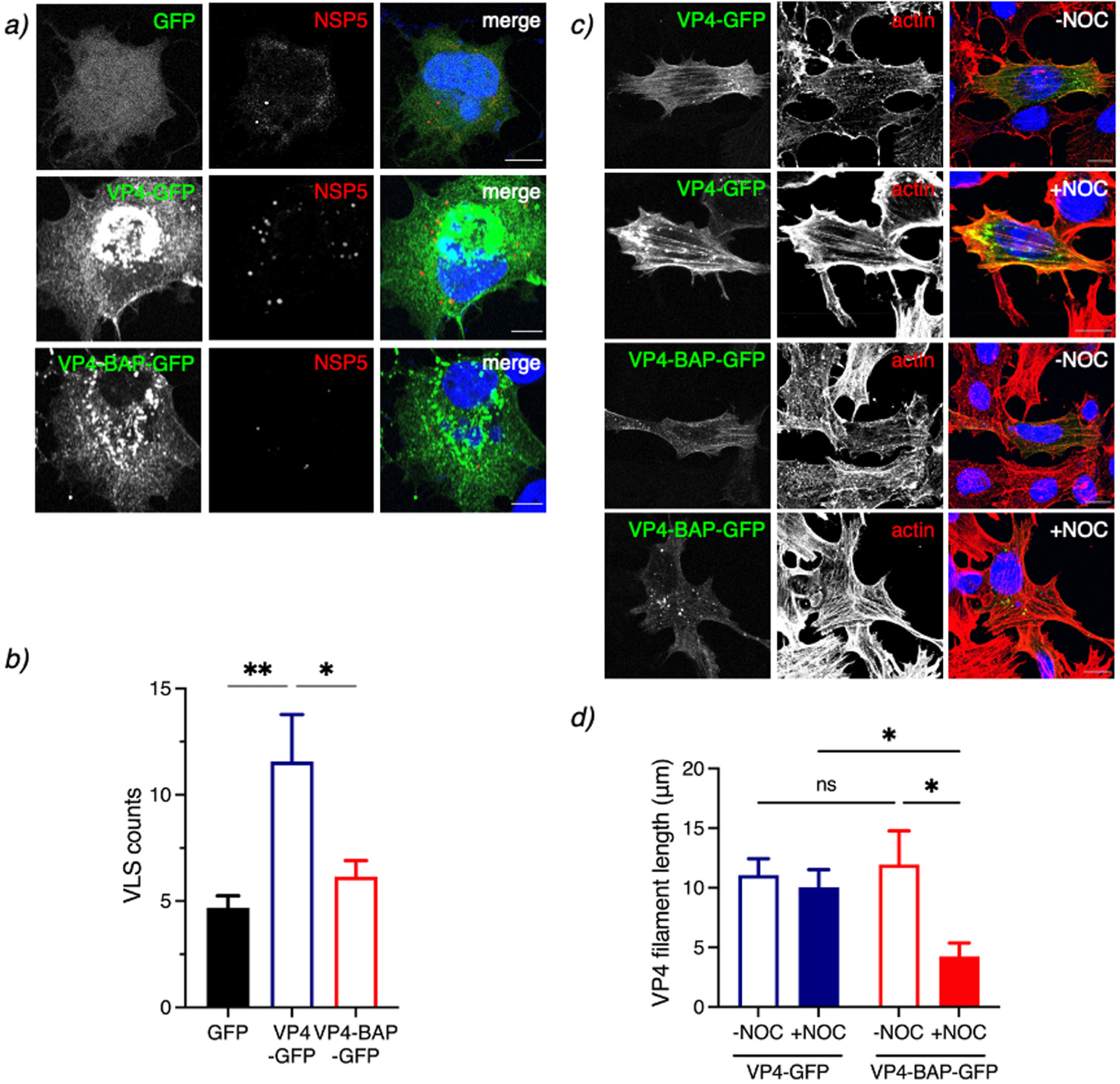
VP4-BAP does not bind to the actin cytoskeleton. **(a)** Immunostaining of MA104 cells showing NSP5 and NSP2 VLSs co-expressed with GFP (top panel), VP4-GFP (middle panel), and VP4-BAP-GFP (bottom panel). Cells were fixed with paraformaldehyde and immunostained to detect VLSs (anti-NSP5, red). Nuclei were stained with DAPI (blue). The scale bar is 10 µm. **(b)** Plot indicating the number of VLSs per cell when co-expressed with GFP, VP4-GFP, and VP4-BAP-GFP.The data represent the mean ± SEM, one-way ANOVA, (*), p<0.05 and (**), p<0.001. **(c)** Immunostaining of MA104 cells expressing VP4-GFP and VP4-BAP-GFP after treatment for 1 h before fixation without (-NOC) or with (+NOC) 10 µM nocodazole. Cells were fixed with methanol at 24 hpt and immunostained for GFP detection (anti-GFP, green) and actin cytoskeleton (anti-actin, red). Nuclei were stained with DAPI (blue). The scale bar is 10 µm. **(d)** Plot for the quantification of VP4-GFP and VP4-BAP-GFP filament lengths associated with actin filaments in cells untreated or treated with nocodazole. The data represent the mean± SD, Welch’s t-test, (****), p<0.0001.

Since VP4 associates with actin cytoskeleton components (45, 47, 48), we hypothesized that VP4-BAP-GFP could have an impaired association with actin filaments. To investigate this possibility, MA104 cells expressing VP4-GFP or VP4-BAP-GFP were untreated or treated with nocodazole for 1h before fixation. Nocodazole treatment induces depolymerization of the microtubule network permitting direct characterization of proteins associated with the actin cytoskeleton. In this context (**Fig 8c**), we stained the cells to detect actin cytoskeleton and noticed that VP4-GFP was associated, as expected, with actin filaments even after nocodazole treatment; while VP4-BAP-GFP was associated with filaments only in the absence of nocodazole. In MT-depolymerized cells, VP4-BAP-GFP formed diffuse small cytosolic aggregates or short filaments. Moreover, while the length of VP4-GFP fibers was comparable in the presence or absence of nocodazole treatment, the VP4-BAP-GFP fibers were significantly shorter in cells treated with nocodazole (**Fig 8d**), suggesting an impaired association of VP4-BAP with actin filaments. Taken together, these results suggest that VP4 promotes the assembly of VLSs mediated by its association with actin filaments.

### Impaired association between the actin cytoskeleton and rRV/VP4-BAP results in delayed viroplasm assembly

RV strain SA11 infection (**Fig 9a** and (25, 61)) reorganizes the actin cytoskeleton, mainly by decreasing the actin stress fiber and redistributing it to the cell cortex. However, this reorganization did not take place in cells infected with rRV/VP4-BAP (**Fig 9b and Fig S3a**) featured by an increment in stress fibers surrounding the viroplasms (**Fig 9b**, yellow open arrows) and contrasting by an actin cell cortex increment in rRV/wt infected cells. Moreover, the actin cytoskeleton is not properly reorganized even in experiments with an increased rRV/VP4-BAP multiplicity of infection (**Fig. S3b**). Interestingly, the MT-network reorganization was attained by both viruses.

**Figure 9.**
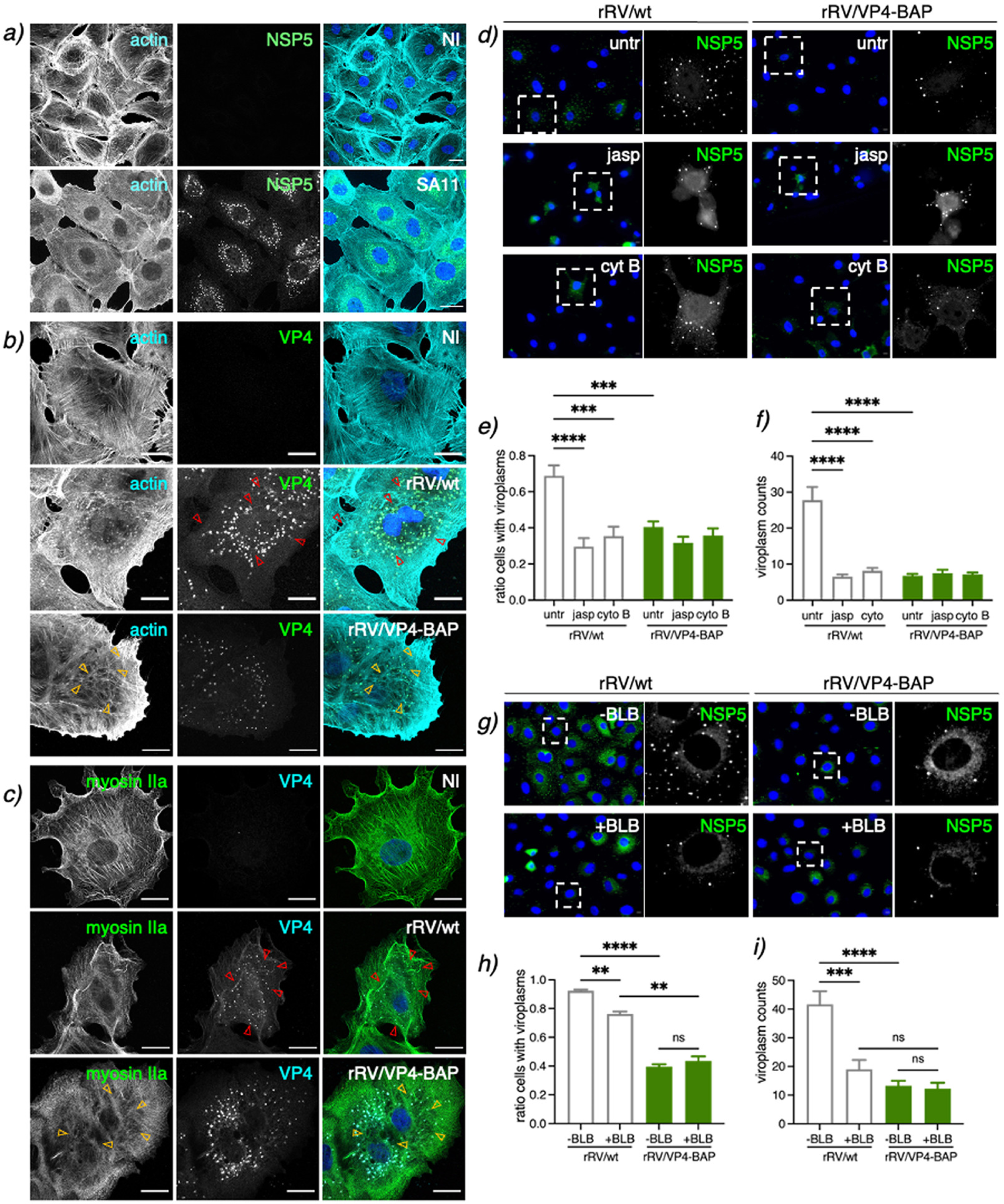
rRV/VP4-BAP impaired association with the actin cytoskeleton. **(a)** Immunostaining of non-infected and SA11-infected MA104 cells. At 6 hpi, cells were fixed with methanol and immunostained to detect viroplasms (anti-NSP5, green) and actin cytoskeleton (anti-actin, cyan). Nuclei were stained with DAPI (blue). The scale bar is 20 µm. Immunostaining of non-infected and rRV/wt or rRV/VP4-BAP-infected MA104 cells. At 6 hpi, cells were fixed with methanol and immunostained for detection of **(b)** VP4 (anti-VP4, green) and actin cytoskeleton (anti-actin, cyan) and **(c)** VP4 (anti-VP5 clone 4G2, cyan) and myosin motor (anti-myosin IIa, green). Nuclei were stained with DAPI (blue). The scale bar is 20 µm. Open yellow and red arrowheads point to stress fibers in the actin cytoskeleton and VP4 fibers, respectively. **(d)** Immunofluorescence of rRV/wt and rRV/VP4-BAP-infected MA104 cells treated at 1 hpi with either 0.5 µM jasplakinolide (jasp; middle panel) or 10 µM cytochalasin B (cyt B; lower panel). At 6 hpi, cells were fixed and immunostained to detect viroplasms (anti-NSP5, green). Nuclei were stained with DAPI (blue). The dashed open box corresponds to the inset of the enlarged image at the right. The scale bar is 10 µm. Plot for the quantification of **(e)** the ratio of cells showing viroplasms and **(f)** the number of viroplasms per cell. **(g)** Immunofluorescence of rRV/wt and rRV/VP4-BAP-infected MA104 cells untreated or treated at 1 hpi with 5 µM blebbistatin (BLB). At 6 hpi, cells were fixed and immunostained to detect viroplasms (anti-NSP5, green). Nuclei were stained with DAPI (blue). The dashed open box is an inset of the enlarged image at the right. The scale bar is 10 µm. Plot for the quantification of **(h)** the ratio of cells showing viroplasms and **(i)** the number of viroplasms per cell.

The cell has a contractile system regulated in part by the reorganization of stress fibers composed by actin and myosin II. We questioned if the molecular motor myosin was also required for viroplasm assembly and if its activity was impaired in rRV/VP4-BAP infected cells. For this purpose, we inspected the localization of paralog myosin IIa in cells infected with either rRV/wt or rRV/VP4-BAP at 6 hpi and compared it with non-infected conditions (**Fig 9c**). In non-infected cells, myosin IIa is homogenously distributed in filaments, stacks, clusters, and continuous structures (62). However, upon rRV/wt infection, continuous myosin structures are lost, and myosin clusters and stacks in the cell cortex are favored (**Fig 9c**, yellow open arrowheads). In contrast, rRV/VP4-BAP infected cells still have continuous structures, mainly in the ventral cell region. Interestingly, VP4 fibrillar morphology is observed in rRV/wt infected cells (**Fig 9b and c**, red open arrows) but not in rRV/VP4-BAP infected cells.

We confirmed the active role of the actin cytoskeleton in the assembly of rRV/wt and rRV/VP4-BAP viroplasms by adding actin inhibitors jasplakinolide (jasp) and cytochalasin B (cyt B) at 1 hpi, a time in which virus internalization and primary virus transcription are well-initiated (**Fig 9d**). In this context, we denoted that rRV/wt viroplasms were sensitive to jasp and cyt B treatment, as the ratio of cells showing viroplasms (**Fig 9e**) and the number of viroplasms per cell (**Fig 9f**) were significantly decreased, reaching the levels observed in rRV/VP4-BAP infected cells in presence or absence of actin inhibitors.

We next inhibited the non-muscular myosin II using blebbistatin (BLB)(63), a small molecule inhibiting both myosin II paralogs a and b (**Fig 9g-i**). Like the actin inhibitors, blebbistatin reduced the ratio of cells with viroplasms (**Fig 9h**) and the number of viroplasms per cell (**Fig 9i**) in cells infected with rRV/wt but not with rRV/VP4-BAP. Our results suggest that viroplasm assembly requires actin and myosin II in a mechanism employing VP4.

### A small peptide mimicking loop K145-G150 rescues the rRV/VP4-BAP phenotype

It has been described that the C-terminal region of VP5* harbors an actin-binding domain (45). We hypothesize that the insertion of a BAP tag in loop K145-G150 interferes with the association of VP5* with the actin cytoskeleton, which results in a delay in viroplasm assembly because of an inability to reorganize the actin cytoskeleton. To prove this hypothesis, we designed a small peptide harboring the amino acid sequence of wt VP4 loop K145-G150 flanked at the N-terminus by an arginine-tail and at the C-terminus by conjugation to fluorescein isothiocyanate (FITC), for peptidic internalization and visualization, respectively. Thus, the sequence of this small peptide corresponds to RRRRRR^143^VV^145^KTTANG^150^SIGQYG^156^-FITC, and was designated <u>s</u>mall <u>p</u>eptide loop K145-G150 (SPL). Of note, SPL was not toxic for cells up to a concentration of 100 µM even after 24 h of treatment (**Fig S4a**). SPL was internalized in cells at 2 h post-treatment and found diffuse in the cytosol **(Fig S4b)**. In the first instance (**Fig 10a**), we expressed VP4-GFP and VP4-BAP-GFP (stained with anti-GFP, red) in BHK-T_7_ cells in the absence or presence of SPL and monitored the filamentous distribution of these proteins. Interestingly, as quantified in **Fig 10b**, the addition of SPL increased the length of VP4-BAP-GFP filaments compared to the untreated sample reaching the same level of the fiber lengths as VP4-GFP. The filamentous distribution of VP4-GFP did not change with SPL treatment. We next interrogated if SPL can improve the replication fitness of rRV/VP4-BAP. For this, SPL was added at 1 or 3 hpi to MA104 cells infected with either rRV/wt or rRV/VP4-BAP and then monitored at 6 hpi for viroplasm formation (**Fig 10c**). Surprisingly, the addition of SPL at 1 hpi improved drastically the number of cells presenting viroplasms in rRV/VP4-BAP-infected cells, reaching similar levels as observed in rRV/wt infected cells (**Fig 10d**). Consistently, the viroplasm size and numbers per cell increased upon SPL treatment at 1 hpi (**Fig 10e and f**). The addition of SPL at 3 hpi resulted in increased viroplasm size but not increased numbers per cell. The treatment of rRV/wt infected cells with SPL did not affect viroplasm size and numbers (**Fig S5c-d**). We then tested the ability of SPL to recover rRV/VP4-BAP virus progeny. For this purpose, rRV/VP4-BAP infected cells were treated at 1 hpi with SPL, and virus progeny was recovered at 12 hpi. As denoted in **Fig 10g**, rRV/VP4-BAP virus progeny formation significantly increased upon treatment with SPL, while the same treatment had no effect on rRV/wt progeny formation. We concluded that SPL boosted the association of VP4-BAP to actin-bundles, thereby rescuing the assembly of viroplasms with a concomitant increase in the production of virus progeny.

**Figure 10.**
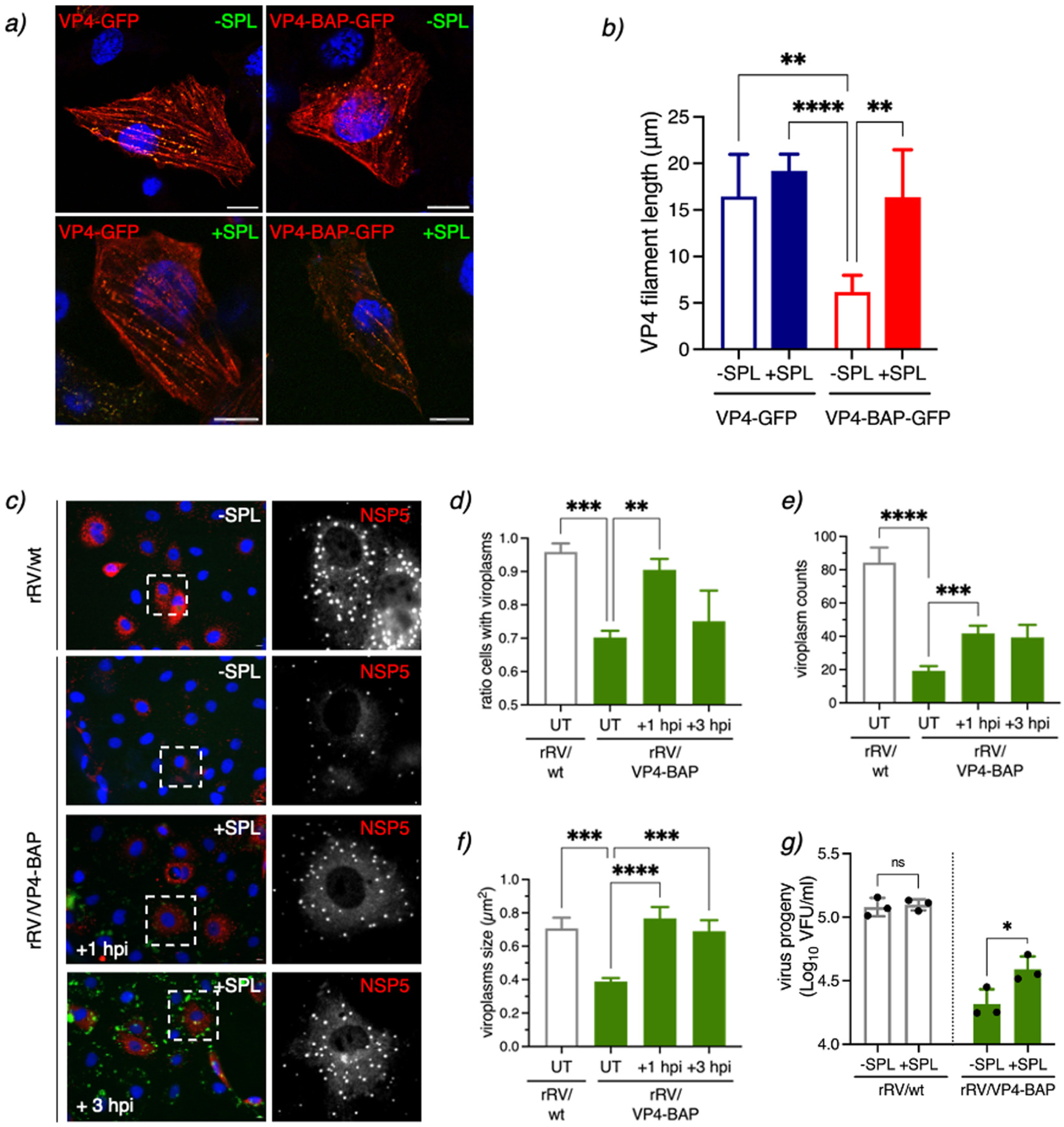
A peptide mimicking loop K145-G150 rescues the rRV/VP4-BAP phenotype. **(a)** Immunofluorescence images at 17 hpt of BHK-T_7_ cells expressing VP4-GFP or VP4-BAP-GFP, untreated or treated with 25 µM SPL added immediately after transfection. Before methanol fixation, cells were treated for 1 h with 10 µM nocodazole. The cells were immunostained for GFP (anti-GFP, red), and nuclei were stained with DAPI (blue). Scale bar is 10 µm. **(b)** Plot quantifying the length of VP4 filaments untreated or treated with SPL as described in **(a)**. Welch ANOVA test, (**), p< 0.01 and (****), p<0.0001. **(c)** Representative immunofluorescence microphotograph of viroplasms of rRV/wt and rRV/VP4-BAP -infected cells untreated (-SPL) or treated (+SPL) at +1 and +3 hpi with 25 µM of SPL, as indicated. Viroplasms were immunostained at 6 hpi with anti-NSP5 (red), and nuclei were stained with DAPI (blue). The dashed white boxes correspond to the enlarged picture in the right columns. Scale bar is 10 µm. Quantification plots for the ratio of cells with viroplasms **(d)** and the numbers **(e)** and size **(f)** of viroplasms per cell of MA104 cells infected with the indicated virus and untreated (UT) or treated with 25 µM SPL at +1 and +3 hpi. The data represent the mean ± SEM using Welch’s ANOVA test, (***), p<0.001 and (****), p<0.0001. **(g)** Plot determining the virus progeny of rRV/wt and rRV/BAP infected cells untreated or treated at 1 hpi with 25 µM SPL. The data represent the mean +SEM of three independent experiments. Welch ANOVA test, (*) p<0.05.

## Discussion

The external coat layer of the RV virion can be modified *in vitro* by adding a specific ratio of VP7 and VP4 proteins to purified DLPs to generate recoated TLPs (rcTLPs) (10-12, 64). This approach provides a valuable tool for studying the VP4 structural requirements for virion internalization (10). However, rcTLPs have methodological limitations and do not allow transferring the parental phenotype to the virus progeny. Moreover, rcTLPs only allow single amino acid substitutions in the VP4 spike protein (65, 66). However, the recently implemented entirely plasmid-based RV reverse genetics technology resulted in a valuable tool for studying several aspects of RV replication and the development of future RV based vaccines because it supports site-directed modifications of RV genome segments of interest. Specifically, RV reverse genetics technology has demonstrated that gs4, encoding VP4, can be artificially reassorted or recombined by transferring human RV strain gs4 into a simian strain SA11 backbone (54).

In the present study, we used an RV reverse genetics system to remodel the spike protein VP4 by incorporating a BAP tag in four independent exposed loops in the VP8* lectin domain. Interestingly, although the four differently modified proteins were efficiently biotinylated in transfected cells, only one of them harboring a BAP tag inserted in the K145-G150 loop, rRV/VP4-BAP, was rescued using RV reverse genetics (54). We hypothesize that those VP4-BAP versions, which were unable to be rescued, could have had strongly compromised their virus progeny because they directly impacted, and as denoted by *in silico* modeling of these structures (**Fig S5a**), *i)* the VP4 transition states from upright (immature) to reverse (mature) (65), *ii)* the VP4 association with specific cellular receptors (36, 37), or *iii)* the incorporation of VP4 in the coat layer.

VP4-BAP and VP4 share similar distribution patterns in infected cells, such as their localization surrounding viroplasms, co-localization in the endoplasmic reticulum with VP7, and incorporation into newly assembled virions. However, we found that rRV/VP4-BAP had a significantly reduced replication fitness compared to rRV/wt. To identify the reason for this, we investigated the various steps involved in RV replication. We show that rRV/VP4-BAP internalization kinetics are comparable to those of the rRV/wt. This internalization kinetics consists of a sequence of events starting with the interaction of the virus with the cell membrane and concluding with the decrease of calcium levels within the endosomes, provoking the loss of the outer layer of the virion, which triggers the release of transcriptionally active DLPs into the cytosol (66-69). We also confirmed that virus transcription and translation are comparable between both viruses. Moreover, there were no differences in rRV/VP4-BAP viroplasm morphology and liquid-like dynamics, as determined by high-resolution electron microscopy and NSP2-mCherry mobility on single-viroplasm FRAP experiments, respectively. However, upon transfection with rRV/VP4-BAP DLPs that bypass the virus physiological internalization pathway, showed a delayed viroplasm formation compared to rRV/wt DLPs, even though NSP5 expression levels were equivalent for both viruses. Collectively, these results indicate that the reduced replication fitness of rRV/VP4-BAP is due to a step between primary virus translation and viroplasm formation. We hypothesize that rRV/VP4-BAP, and specifically VP4-BAP, was defective in associating with the actin cytoskeleton because we observed *i)* a delayed recovery of rRV/VP4-BAP viroplasms after 1,6-HD treatment, *ii)* inability of VP4-BAP-GFP to increase the VLS counts, *iii)* absence of co-localization of VP4-BAP-GFP with actin stress fibers, *iv)* inability of rRV/VP4-BAP to reorganize actin cytoskeleton, and *v)* insensitivity of rRV/VP4-BAP viroplasms for drugs depolymerizing actin filaments and inhibiting myosin II molecular motor. In contrast, rRV/wt infection led to the reorganization of the actin cytoskeleton, and rRV/wt viroplasms were sensitive to actin-depolymerizing and myosin inhibitor drugs linking actin with viroplasm assembly. Additionally, the numbers of VLS duplicated when co-expressed with VP4-GFP. Previous studies (45, 49) demonstrated the ability of VP4 to associate with actin in the absence of other virus proteins, mainly through an actin-binding domain present in the C-terminal region of VP5* (residues 713-776) when in cooperation with the coiled-coil domain (residues 481-574). In this context, we also present evidence that the sole expression of VP4 allows its association with actin. But importantly, by employing rRV/VP4-BAP infection, we show that the association of VP4 with actin is essential to catalyze the formation of viroplasms. This finding provides an additional function to VP4 in RV replication.

Interestingly, incorporating a small peptide mimicking loop K145-G150 of VP4 during rRV/VP4-BAP infection reverted the impaired formation of viroplasm with a significant improvement in virus progeny production. We concluded that rRV/VP4-BAP replication is enhanced because SPL raised the association of actin with VP4-BAP during virus infection and because we observed that VP4-BAP, in the absence of other viral proteins, associated with actin filament when expressed with SPL. We, therefore, hypothesize that the VP8* subunit is involved in at least one of these three aspects that render the viroplasms assembled and stabilized: *i)* association of VP8* with a yet undescribed host component, *ii)* a reorganization of VP5*-VP8* association or *iii)* a direct role of VP8* over another RV protein in viroplasm assembly. The role of the actin cytoskeleton in the RV life cycle, particularly internalization and egress, has been previously demonstrated (39, 47, 70). Here, we show that the actin cytoskeleton is also involved in viroplasm formation, an essential intermediate stage of RV replication. It has been described that the silencing of Rac1, a member of the Rho family of small GTPases playing a major role in actin- and microtubule-cytoskeleton dynamics, leads to a decrease in RV progeny formation in a process downstream of cell entry (70). We therefore cannot exclude activation of Rac1 by VP4, allowing reorganization of the actin cytoskeleton for an efficient assembly of the viroplasms.

Finally, specific *in vivo* biotinylation of cellular targets can be achieved by adding a BAP tag to the protein of interest and co-expressed with the *E.coli*-derived biotin ligase, BirA (71). This method is a powerful tool for versatile applications, such as identifying highly complex interactomes (72, 73), and permits batch protein and subviral particle (51, 52) refinement at high purity and in physiological conditions. The incorporation of a BAP tag in RV VP6, allowed the preparation and purification of replication-competent DLPs (52). Identifying a permissive target site in loop K145-G150 of VP4 spike protein for the insertion of an exogenous peptide may impact the RV field. This VP4 modification favors the insertion of peptides required for super-resolution microscopy or DNA-paint technologies (e.g., Halo or BC2 tags) to dissect debated aspects of RV entry. In addition, this VP4 modification technology could permit the incorporation of antigenic peptides for vaccine development. Although it is well-known that the current oral RV vaccines elicit an efficient immune response (74, 75), rRV harboring a modified VP4 could provide an improved vaccination platform for the display of other antigens fostering the development of a new generation of dual vaccines.

## Material and Methods

### Cells and viruses

MA104 cells (embryonic African green monkey kidney cells; ATCC CRL-2378) were grown in Dulbecco’s modified Eagle’s medium (DMEM) (Life Technologies) containing 10% fetal calf serum (FCS) (AMIMED; BioConcept, Switzerland) and penicillin (100 U/ml)–streptomycin (100 μg/ml) (Gibco, Life Technologies). MA/cytBirA and MA/NSP2-mCherry (59) cell lines were grown in DMEM supplemented with 10% FCS, penicillin (100 U/ml)-streptomycin (100µg/ml) and 5µg/ml puromycin (InvivoGen, France). BHK-T_7/9_ (baby hamster kidney stably expressing T_7_ RNA polymerase) cells were kindly provided by Naoto Ito (Gifu University, Japan)(76) and cultured in Glasgow medium supplemented with 5% FCS, 10% tryptose phosphate broth (Sigma-Aldrich), 10% FCS, penicillin(100 U/ml)-streptomycin (100µg/ml), 2% nonessential amino acids and 1% glutamine.

rRV/wt (59), rRV/VP4-BAP, and simian rotavirus strain SA11 (G3P6[1])(77) were propagated, grown, and purified as previously described (78). Virus titer was determined as viroplasm forming units per ml (VFU/ml) as described by Eichwald et al. 2012 (25). The T_7_ RNA polymerase recombinant vaccinia virus (strain vvT7.3) was amplified as previously described (79).

### Cell line generation

MA/cyt-BirA cell line was generated using the PiggyBac technology (80). Briefly, 1x10^5^ MA104 cells were transfected with the pCMV-HyPBase (80) and transposon plasmids pPB-cytBirA using a ratio of 1:2.5 with Lipofectamine™ 3000 transfection reagent (Invitrogen, Thermo Fisher Scientific) according to the manufacturer’s instructions. The cells were maintained in DMEM supplemented with 10% FCS for three days and then selected for four days in DMEM supplemented with 10% FCS and 5 μg/ml puromycin (59).

### Reverse genetics

rRV/VP4-BAP was prepared as described previously (59, 81) using a pT_7_-VP4-BAP instead of pT_7_-VP4. Briefly, monolayers of BHK-T_7_ cells (4×10^5^) cultured in 12-well plates were co-transfected using 2.5 μl of TransIT-LT1 transfection reagent (Mirus) per microgram of DNA plasmid. The mixture comprised 0.8 μg of SA11 rescue plasmids: pT_7_-VP1, pT_7_-VP2, pT_7_-VP3, pT_7_-VP4-BAP, pT_7_-VP6, pT_7_-VP7, pT_7_-NSP1, pT_7_-NSP3, pT_7_-NSP4, and 2.4 μg of pT_7_-NSP2 and pT_7_-NSP5 (82, 83). Additionally, 0.8 μg of pcDNA3-NSP2 and 0.8 μg of pcDNA3-NSP5, encoding NSP2 and NSP5 proteins, were co-transfected to increase rescue efficiency (59, 81). Next, cells were co-cultured with MA104 cells for three days in serum-free DMEM supplemented with trypsin from porcine pancreas (0.5 μg/ml final concentration) (T0303-Sigma Aldrich) and lysed by freeze-thawing. Then, 300 μl of the lysate was transferred to new MA104 cells and cultured at 37°C for four days in serum-free DMEM supplemented with 0.5 μg/ml trypsin until a visible cytopathic effect was observed. The modified genome segments of rescued recombinant rotaviruses were confirmed by specific PCR segment amplification followed by sequencing (59).

### Antibodies and Chemicals

Guinea pig anti-NSP5, guinea pig anti-RV, goat anti-RV, mouse monoclonal anti-NSP5 (clone 2D2), and rabbit anti-VP4 were described previously (25, 84-87). Mouse monoclonal anti-VP5 (clone 2G4) and mouse monoclonal anti-VP7 (clone 159) was kindly provided by Harry Greenberg (Stanford University, CA, USA). Rabbit anti-simian rotavirus VP4 was purchased from Abcam. Mouse mAb anti-glyceraldehyde dehydrogenase (GAPDH) (clone GAPDH-71.1), mouse anti-alpha tubulin (clone B-5-1-12), and mouse mAb anti-β-actin (clone AC-74) were purchased to Merck. Mouse mAb anti-GFP clone C-2) was purchased from Santa Cruz Biotechnology, Inc. Mouse mAb anti-α-tubulin was directly conjugated to Atto 488 using lightning-link™Atto 488 conjugation kit from Innova Bioscience, UK. Streptavidin-HRP was purchased from Merck. Streptavidin-Alexa 555 and secondary antibodies conjugated to Alexa 488, Alexa 594, Alexa 647, Alexa 700 (ThermoFisher Scientific).

1,6-hexanediol, nocodazole, jasplakinolide, and cytochalasin B were purchased from Merck. (-)-Blesbbistatin was purchased from Cayman Chemical, USA.

Amino acids and their derivatives were purchased from Advanced ChemTech, Novabiochem, Iris Biotech GMBH, Sigma-Aldrich, PolyPeptide, Space peptides and GL BioChem. Amino acids were used as the following derivatives Fmoc-Arg(Pbf)-OH, Fmoc-Leu-OH, Fmoc-Lys(Boc)-OH, Fmoc-Lys(Fmoc)-OH, Fmoc-Pro-OH, Fmoc-Val-OH, Fmoc-Trp(Boc)-OH, Fmoc-Ile-OH, Fmoc-Gln(Trt)-OH, Fmoc-Tyr(tBu)-OH, Fmoc-Ala-OH and Rink Amide AM resin (loading: 0.38 mmol·g^−1^) and were purchased from Sigma Aldrich. OxymaPure (hydroxyiminocyanoacetic acid ethyl ester) and DIC (*N*,*N*^’^-diisopropyl carbodiimide) were purchased from Iris Biotech GMBH. 5(6)-carboxyfluorescein (CF) was from Sigma. EM104 10ml glass syringes from Sanitex international.

### DNA plasmids

pcDNA-VP4-SA11 was obtained by RT-PCR amplification of VP4 ORF of gs4 of simian rotavirus strain SA11 (88) using specific primers to insert *Hind*III and *Xho*I sites, followed by ligation into those sites in pcDNA3 (Invitrogen). pcDNA-VP4-*Kpn*I/*BamH*I was built by insertion of point mutations in pcDNA-VP4-SA11 using the QuikChange site-directed mutagenesis kit and protocol (Agilent) to insert *Kpn*I and *BamH*I restriction sites in VP4. pcDNA-VP4-BAP (blue), (orange), (pink), and (green) were obtained by ligation between *Kpn*I and *BamH*I of pcDNA-VP4-*Kpn*I/*BamH*I, a synthetic DNA fragment (GenScript®) containing BAP tag in VP4 loops in amino acid regions 96-101, 109-114, 132-137 and 145-150, respectively. The BAP tags are flanked by *BspE*I and *Nhe*I restriction sites for easy tag replacement.

RV plasmids pT_7_-VP1-SA11, pT_7_-VP2-SA11, pT_7_-VP3-SA11, pT_7_-VP4-SA11, pT_7_-VP6-SA11, pT_7_-VP7-SA11, pT_7_-NSP1-SA11, pT_7_-NSP2-SA11, pT_7_-NSP3-SA11, pT_7_-NSP4-SA11, and pT_7_-NSP5-SA11 were previously described (82). pcDNA3-NSP5 and pcDNA3-NSP2 were already described (59). pT_7_-VP4-BAP (blue), (orange), (green), and (pink) were obtained by inserting a synthetic DNA fragment (Genscript) encoding for the VP4 protein-encoding BAP tag flanked by *Mfe*I and *Nde*I restriction enzymes sites and ligated into those sites in the pT_7_-VP4-SA11.

pPB-cytBirA was obtained from a synthetic DNA fragment (Genscript) containing the BirA enzyme open reading frame of *Escherichia coli* (UniProt accession number: P06709) and inserted in the pPB-MCS vector (81) using *Nhe*I-*BamH*I restriction enzymes sites.

pcDNA-NSP5(SA11) and pcDNA-NSP2(SA11) were previously described (24, 89). pCI-VP4-GFP and pCI-VP4-BAP plasmids were obtained from PCR amplification of pT_7_-VP4-SA11 and pT_7_-VP4-BAP (green) using specific primers to insert *Nhe*I and *Mlu*I sites, followed by ligation in-frame on those sites in pCI-GFP. Thus, the GFP fragment was PCR amplified from pEGFP-N1 (Clontech) using specific primers to insert *Mlu*I/*Not*I restriction enzyme sites and ligated on those restriction enzyme sites into pCI-Neo (Promega). All the oligonucleotides were obtained from Microsynth AG, Switzerland. A list of all DNA sequences synthesized is provided in **Table S1**.

### Streptavidin-supershift assay and immunoblotting

The assay was performed as described by Predonzani et al (51). Briefly, cell extracts were lysed in TNN lysis buffer (100mM Tri-HCl pH8.0, 250 mM NaCl, 0.5% NP-40, and cOmplete protease inhibitor (Roche)) and centrifuged for 7 min at 15’000 rpm and 4°C. The supernatant was exhaustively dialyzed against PBS (phosphate-buffered saline, 137 mM NaCl, 2.7 mM KCl, 8 mM Na_2_HPO_4_, and 2 mM KH_2_PO_4_ pH 7.2) at 4°C and heated for 5 min at 95°C in Laemmli sample buffer. Samples were incubated for 1 h at 4°C with 1 µg streptavidin (Sigma) and then resolved in SDS-polyacrylamide gel under reducing conditions. Proteins were transferred to nitrocellulose 0.45 µm (90) and incubated with corresponding primary and secondary antibodies. Secondary antibodies were conjugated to IRDye680RD or IRDye800RD (LI-COR, Germany) for protein detection and quantification in Odyssey® Fc (LI-COR Biosciences).

### Virus fitness curve

The experiment was performed as described previously (91) with some modifications. MA104 cells (1×10^5^) seeded in 24-well plates were infected with rRV at an MOI of 10 VFU/cell. The virus was allowed to adsorb for 1 h at 4°C, followed by incubation at 37°C in 500 μl DMEM. At the indicated time points, the plates were frozen at −80°C. Each time point was performed in triplicated. The cells were freeze-thawed for three cycles, harvested, and centrifuged at 900 × *g* for 5 min at 4°C. The supernatant was recovered and activated with 80 μg/ml of trypsin for 30 min at 37°C. Two-fold serial dilutions were prepared and used to determine the viral titers described previously (59, 81).

### Fluorescence labeling of purified rRV

100 µl of purified and trypsin activated biotinylated rRV/VP4-BAP was incubated with 1 µl of streptavidin-Alexa Fluor 555 (2mg/ml) (ThermoFisher Scientific) for 1 h at room temperature. The tube was snaped every 20 min. Unbound streptavidin and streptavidin conjugated to rRV/VP4-BAP were separated by loading the 50 µl reaction mixture on top of 100 µl of a 20% sucrose-PBS cushion followed by centrifugation for 40 min at 20 psi in an air-driven ultracentrifuge (Airfuge, Beckman Coulter). Pellet was resuspended in 20 µl TBS (25 mM Tris-HCl, pH 7.4, 137 mM NaCl, 5 mM KCl, 1 mM MgCl_2_, 0.7 mM CaCl_2_, 0.7 mM Na_2_HPO_4_, 5.5 mM dextrose).

### Immunofluorescence

For virus internalization experiments, 1 µl of rRV particles conjugated to StAV-Alexa555 diluted in 50 µl of DMEM was adsorbed over MA104 cells for 15 min and kept on a metal tray cooled to -20°C. Cells were then transferred to 37°C and fixed at the indicated time-points with ice-cold methanol for 3 min on dry ice.

For later times post-infection, the virus was adsorbed for 1 h at 4°C in a reduced volume. Then, cells were transferred to 37°C and treated at the indicated time points with 100 µM biotin in serum-free DMEM. Cells were fixed when indicated in 2% paraformaldehyde in phosphate-buffered saline (PBS) for 10 min at room temperature or in ice cold-methanol for 3 min at -20°C.

VLS experiments were performed as described by Buttafuoco *et al*. (90). In experiments using the inhibitors, the drug was added at 1 hpi and maintained until 6 hpi. The concentrations used 0.5 µM jasplakinolide (61, 70), 10 µM cytochalasin B (92) and 5 µM blebbistatin (63) were described elsewhere.

All immunofluorescence assays were processed as described by Buttafuoco *et al*. (90). Images were acquired using a confocal laser scanning microscope (CLSM) (DM550Q; Leica). Data were analyzed with the Leica Application Suite (Mannheim, Germany) and Image J (93).

### LLPS assay

MA/NSP2-mCherry cells were seeded at a density of 1.2 x 10^4^ cells per well 8-wells Lab-Tek® Chamber Slide™ (Nunc, Inc. Cat #177402). For RV infection, the virus was adsorbed at MOI of 25 VFU/cell diluted in 30 µl of serum-free DMEM, incubated at 4°C for 1 h in an orbital shaker, and then volume filled to 100µl with serum-free DMEM followed by incubation at 37°C. At 5 hpi, the medium was replaced by medium containing 3.5% 1,6-hexanediol in 2% FCS-DMEM and cells were incubated for 6 min at 37°C. Then the drug was washed out by removing the medium, washing the cells three times with PBS, adding fresh 2% FCS-DMEM, and incubating at 37°C. At the designated time post-recovery, cells were fixed with 2% PFA for 10 min at room temperature. Finally, nuclei were stained by incubating cells with 1 µg/ml of DAPI (4’,6-diamidino-2-phenylindole) in PBS for 15 min at room temperature. Samples were mounted in ProLong™ Gold antifade mountant (ThermoFisher Scientific), and images were acquired using a fluorescence microscope (DMI6000B, Leica). Data were analyzed using ImageJ software (version 2.1.0/1.53; https://imagej.net/Fiji).

### Quantification of viroplasms

The number of viroplasms was acquired and analyzed as previously described (25, 61, 92). Data analysis was performed using Microsoft® Excel for Mac version 16.58. Statistical analysis, unpaired parametric Welch’s t-test comparison post-test, and plots were performed using Prism 9 for macOS version 9.3.1 (GraphPad Software, LLC).

### Rotavirus electropherotype

Rotavirus genome extraction and visualization were performed as previously described (94). Briefly, MA104 cells at a density of 3 x 10^5^ cells per well in a 6-well multiwell plate were infected with a virus at MOI of 10 VFU/cell and incubated in 1 ml serum free DMEM until complete cytopathic effect was reached. The cells and supernatant were harvested, followed by three cycles of liquid nitrogen freeze and 37°C water bath. Then, the samples were mixed vigorously at a ratio of 1:1 with saturated phenol solution pH 4.3 (Merck) and centrifuged for 15 min at 13’000 rpm. The aqueous phase was recovered, and the previous step was repeated. Then, the RNA in the recovered aqueous phase was precipitated by mixing with 0.1 vol 3M Na Acetate at pH 5.2 and 2 vol of 100% ethanol. Samples were incubated for 30 min at -80°C and then centrifuged at 13’000 rpm for 30 min and 4°C. The pellet was resuspended in 15 µl distilled water and mixed with 10 µl Gel Loading dye 6X (New England BioLabs). The samples were migrated in an 8.5% SDS-polyacrylamide gel at 180 Volts for 120 min, followed by staining with GelRed® Acid gel Stain (Biotium) for 30 min. Images were acquired at Odyssey® FC (LI-COR Biosciences).

### Transmission electron microscopy

MA104 cells were seeded at 1 × 10^5^ cells in a 2-cm^2^ well onto sapphire discs and infected with either rRV/wt or rRV/VP4-BAP at an MOI of 50 VFU/ml. At 6 and 12 hpi, the sapphire discs were collected, fixed with 2.5% glutaraldehyde in 100 mM Na/K-phosphate buffer, pH 7.4, for 1 h at 4°C and then kept in 100 mM Na/K-phosphate buffer overnight at 4°C. Afterward, samples were postfixed in 1% osmium tetroxide in 100 mM Na/K-phosphate buffer for 1 h at 4°C, dehydrated in a graded ethanol series starting at 70%, followed by two changes in acetone, and embedded in Epon. Ultrathin sections (60 to 80 nm) were cut and stained with uranyl acetate and lead citrate.

For staining biotinylated TLPs with streptavidin-gold, purified particles were dialyzed overnight at 4°C in TNC buffer (10 mM Tris-HCl, pH 7.5, 140 mM NaCl, 10mM CaCl_2_). The TLPs were adsorbed for 10 min on carbon-coated Parlodion films mounted on 300-mesh copper grids (EMS). Samples were washed once with water, fixed in 2.5% glutaraldehyde in 100 mM Na/K-phosphate buffer, pH 7.0, for 10 min at room temperature, and washed twice with PBS before incubation for 2 h at room temperature with 10 µl streptavidin conjugated to 10 nm colloidal gold (Sigma-Aldrich, Inc). Before use, the streptavidin-gold conjugate was treated as described previously to separate unconjugated streptavidin from streptavidin-conjugated to colloidal gold (95). The viral particles were washed three times with water and stained with 2% phosphotungstate, pH 7.0, for 1 min at room temperature. Samples were analyzed in a transmission electron microscope (CM12; Philips, Eindhoven, The Netherlands) equipped with coupled device (CCD) cameras (Ultrascan 1000 and Orius SC1000A; Gatan, Pleasanton, CA, USA) at an acceleration voltage of 100 kV.

For calculation of the diameter of virus particles by negative staining, the area of each virus particle was calculated using Imaris software (version 2.1.0/1.53c; Creative Commons license) and then converted to the diameter as follows: 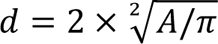, where A is the area and d is the diameter of the particle, respectively.

### siRNA reverse transfection

For silencing gs4 of SA11 strain, the following siRNA pool: siVP4-25 (5’-uugcucacgaauucuuauatt-3’), siVP4-931 (5’-gaaguuaccgcacauacuatt-3’) and siVP4-1534 (5’-auugcaaugucgcaguuaatt-3’) was designed and synthesized by Microsynth AG (Switzerland). siRNA-A (sc-37007, Santa Cruz Biotechnology) was used as negative siRNA control. siRNA reverse transfection was performed by mixing 1.2 µl siRNA 5 µM with 1 µl lipofectamine RNAiMAX transfection reagent (Invitrogen, ThermoFisher Scientific) to a final volume of 100 µl with Opti-MEM® (Gibco, ThermoFisher Scientific) in a well of 24-well plate and incubated for 20 min at room temperature. To reach a 10 nM siRNA final concentration, 2 x 10^4^ cells diluted in 500 µl DMEM supplemented with 10% FCS were added on top. Samples were incubated for 60 h prior to analysis. Thus, cells were infected with simian RV strain SA11 at MOI 12 VFU/cell as described previously (25, 90, 91).

### FRAP

1.2 x 10^4^ MA/NSP2 cells per well were seeded in µ-Slide 18-well glass-bottom plates (Ibidi). Cells were RV-infected at MOI of 15 VFU/cell and kept in DMEM-SF. At 4.5 hpi, the cells were counterstained with Hoechst 33342 diluted in FluoroBRITE DMEM (Gibco, Cat.No. A18967-01) at a concentration of 1 µg/ml, incubated for 30 min at 37°C, and subjected to FRAP analysis. FRAP experiments were performed with an SP8 Falcon confocal laser scanning microscope (CLSM) from Leica equipped with a 63x objective (NA 1.4) using the FRAP function of the LasX software (Leica) as follows: a circular area of 2 µm in diameter, encompassing an entire viroplasm, was bleached with the 405 nm and 481 nm lasers simultaneously, each at 100% laser power, for 20 iterations. The fluorescent recovery was monitored by taking fluorescence images of the mCherry channel every 2 seconds for 140 seconds. For each FRAP acquisition, a circular area of 2 µm, encompassing an entire unbleached viroplasm in the same cell, was used as the fluorescent control, and a squared area (5 µm x 5 µm) outside of a cell was chosen as the background. The entire FRAP dataset was analyzed with MatLab (MATLAB R2020b, Mathworks) using the FRAP-tool source code from easyFRAP (Cell Cycle Lab, Medical School, University of Patras). Fully normalized data were used to generate FRAP diagrams and calculate the recovery half-times (T-half) and mobile fractions from independent measurements. Representative images were taken and processed for each FRAP experiment using the Imaris software v9.5 (Bitplane, Oxford Instruments). Fluorescent intensities of FRAP movies were normalized using a customized Fiji pipeline (93).

### Quantitative RT-PCR

MA104 cells at a density of 5 x 10^5^ cells per well in 6 well multiwell plates were RV-infected at an MOI of 18 VFU/cell. The virus was adsorbed for 1 h at 4°C. At 4 hpi, RNA was extracted using a quick RNA miniprepPlus kit (Zymo Research) according to the manufacturer’s instructions. cDNA synthesis was prepared with 1 µg of RNA using an AMV reverse transcription system (Promega) and random primers according to the manufacturer’s instructions. Then 2 µl of 1:10 diluted cDNA was mixed with 0.25 µl forward primer (10 pmol/µl), 0.25 µl reverse primer (10 pmol/µl) (**Supplementary Table 2**), 10 µl SYBR™ Green PCR master mix (Applied biosystems), and 7.5 µl nuclease-free water followed by incubation at QuantStudio3 (Applied Biosystems, ThermoFisher Scientific) using standard amplification protocol with an annealing temperature of 60 °C. The relative expression of genes was calculated with the formula 2^-ΔCt^, where ΔCt = Ct target gene-Ct endogenous control gene. HPRT-1, SDHA and GAPDH were used as an endogenous control housekeeping gene. Data were analyzed using Microsoft Excel for MAC (version 16.58).

Statistical analysis and plots were done using Prism9 for macOS (version 9.3.1) (GraphPad Software, LLC).

### Purification and transfection of DLPs

RV amount sufficient to infect 1 x 10^6^ cells at MOI of 15 VFU/cell was diluted up to 110 µl with TBS. Then EDTA pH 8.0 was added to a final concentration of 10 mM. The samples were incubated for 1 h at 37°C with gentle mixing and centrifuged at 3000 rpm for 2 min. The supernatant was recovered, loaded on top of a 100 µl cushion composed of 20% sucrose in PBS, and ultracentrifuged at 20 psi for 60 min on an air-driven ultracentrifuge (Airfuge, Beckman Coulter). For quality control, DLPs were monitored by negative staining electron microscopy. The DLPs were normalized using Pierce™ Coomassie protein assay kit (ThermoFisher Scientific) to determine the amount of total protein followed by an indirect ELISA to normalize to rotavirus protein using as primary antibody a guinea pig anti-rotavirus, which detected mainly VP6. Thus, DLPs were transfected by diluting them in 12.5 µl Opti-MEM and mixed with 0.75 µl Lipofectamine 2000 (Invitrogen) in 12.5 µl Opti-MEM. The mixture was incubated for 20 min at room temperature and added onto 1 x 10^4^ MA104 cells per well in a 96-well black wall tissue culture plate (Greiner). At 6 hpi, cells were fixed with paraformaldehyde and prepared for immunofluorescence as described above. Images were acquired using a CLSM and then processed with Image J2 version 2.3.0/1.53f.

### In-cell western assay

RV-infected MA104 cells (1 x 10^4^) were seeded in a 96-well black tissue culture plate (Greiner). At indicated times post-infection, cells were fixed with 2% paraformaldehyde in PBS for 10 min at room temperature, followed by permeabilization with 0.1% triton X-100-PBS for 10 min at room temperature. Cells were blocked with 2% BSA in PBS for 1 h at room temperature and then incubated with primary antibodies diluted in blocking buffer for 1 h at room temperature in the shaker. The cells were incubated with the corresponding secondary antibody conjugated to IRDye® 800 CW (LI-COR). The cell signal was normalized using CellTag™ 700 stain (LI-COR) diluted at 1:800. The cells were washed three times with 0.01% Tween 20 in PBS between incubations. Samples were acquired using an Odyssey CLx (LI-COR) followed by data analysis and normalization in Microsoft® Excel for Mac version 16.58. Statistical analysis, unpaired Student’s t-test, and plots were performed using Prism 9 for macOS version 9.3.1 (GraphPad Software, LLC).

### Virus attachment assay

Rotavirus binding was determined by a nonradioactive binding assay as described previously in detail by Zarate *et al.* (56). The plates were coated with goat anti-rotavirus (diluted 1:5000) to capture the virus. In addition, guinea pig anti-rotavirus (diluted 1:5000) or streptavidin-HRP (diluted 1:500, Sigma) were used to detect the virus. The reaction was developed using 100 µl of Pierce TMB substrate kit (ThermoFisher) and stopped with 100 µl 1M H_2_SO_4_. Samples were recorded at an absorbance of 450 nm using an Infinite M Plex (Tecan) plate reader. Data analysis was performed using Microsoft® Excel for Mac version 16.58. Statistical analysis, semilog line nonlinear regression, and plots were performed using Prism 9 for macOS version 9.3.1 (GraphPad Software, LLC).

### Linear peptide synthesis

The linear peptide was synthesized using an automated peptide synthesis system developed in our laboratory. In here, 200 mg of resin was swelled in 6 ml 100% dimethylformamide (DMF) for 5 min at 60 °C with nitrogen bubbling using a Teflon tube of 1/8-inch inner diameter. Importantly, the bubbling with nitrogen is kept during the entire synthesis process. The Fmoc-protecting groups of the resin were removed by washing twice with 6 ml solution of 20 % piperidine in DMF. For the first deprotection, the sample was incubated for 1 min at 60 °C followed by aspiration. The second deprotection was obtained by incubating for 4 min at 60 °C followed by four washes with 7 ml DMF. The coupling of amino acids to unprotected resin or to the subsequent unprotected amino acids in the elongated peptide coupled to the resin was performed by adding twice 7.5 ml mixture of respective amino acid (5 eq/amine – 3 ml of 200mM), OxymaPure (7.5 eq./amine – 2ml of) and DIC (10 eq./amine – 2.5ml of) dissolved in DMF. The amino acids were coupled for two times 8 min at 60 °C with two times with 7 ml DMF wash after the first coupling and three times with 7 ml DMF wash after the second coupling. After each coupling, the newly bound amino acid was deprotected as mentioned above. The coupling reaction time was specifically extended to 10 min for Asp, Glu and Arg and temperature was lowered to 50 °C for Asp and Glu. The final deprotection was performed in the same manner as explained above. The newly synthesized peptide was washed twice with methanol and dried under vacuum. The peptide cleavage was carried out using TFA/TIS/H2O (94:5:1 v/v/v) for 4 h in 10 ml plastic syringe mounted on a rotter. Then, the peptide was collected in 50 ml tubes, initially with gravity flow and subsequently, by inserting the plunger into the syringe. After filtration, the peptide was precipitated with 50 ml ice cold tert-butylmethylether (TBME), centrifuged at 4400 rpm for 15 min, and washed twice with TBME. For purification of the crude peptide, it was dissolved in 10 ml of a solution containing100% mQ-H_2_O, 0.1% TFA, subjected to preparative RP-HPLC and obtained as TFA salt after lyophilisation. MS spectra were provided by Mass Spectrometry of the Department of Chemistry and Biochemistry at the University of Bern.

## Acknowledgments

We thank Jakub Kubacki for his support of deep sequencing technology. This work has been supported by the University of Zurich. A pre-doctoral ICGEB fellowship also supported this project for GP and GDL. KG was supported by Diaconis-AMM Berner Stellennetz, Switzerland.

The authors declare no conflict of interest.

